# Contrasting paternal and maternal genetic histories of Thai and Lao populations

**DOI:** 10.1101/509752

**Authors:** Wibhu Kutanan, Jatupol Kampuansai, Metawee Srikummool, Andrea Brunelli, Silvia Ghirotto, Leonardo Arias, Enrico Macholdt, Alexander Hübner, Roland Schröder, Mark Stoneking

## Abstract

The human demographic history of Mainland Southeast Asia (MSEA) has not been well-studied; in particular there have been very few sequence-based studies of variation in the male-specific portions of the Y chromosome (MSY). Here, we report new MSY sequences of ∼2.3 mB from 914 males, and combine these with previous data for a total of 928 MSY sequences belonging to 59 populations from Thailand and Laos who speak languages belonging to three major MSEA families: Austroasiatic (AA), Tai-Kadai (TK) and Sino-Tibetan (ST). Among the 92 MSY haplogroups, two main MSY lineages (O1b1a1a* (O-M95*) and O2a* (O-M324*)) contribute substantially to the paternal genetic makeup of Thailand and Laos. We also analyse complete mtDNA genome sequences published previously from the same groups, and find contrasting pattern of male and female genetic variation and demographic expansions, especially for the hill tribes, Mon, and some major Thai groups. In particular, we detect an effect of post-marital residence pattern on genetic diversity in patrilocal vs. matrilocal groups. Additionally, both male and female demographic expansions were observed during the early Mesolithic (∼10 kya), with two later major male-specific expansions during the Neolithic period (∼4-5 kya) and the Bronze/Iron Age (∼2.0-2.5 kya). These two later expansions are characteristic of the modern AA and TK groups, respectively, consistent with recent ancient DNA studies. We simulate MSY data based on three demographic models (continuous migration, demic diffusion and cultural diffusion) of major Thai groups and find different results from mtDNA simulations, supporting contrasting male and female genetic histories.

## Introduction

Thailand and Laos occupy a key location in the center of Mainland Southeast Asia (MSEA; Figure 1), which is undoubtedly one of the factors facilitating the extensive ethnolinguistic diversity there, as there are 68 recognized groups in Thailand and 82 groups in Laos, belonging to five language families (Simons and Fennig 2018). The prehistoric peopling of the area of present-day Thailand and Laos has been documented by several archaeological studies (Shoocongdej 2006; Demeter et al. 2012; Higham 2014; Higham 2017) and investigated further by recent ancient DNA studies (Lipson et al., 2018; McColl et al., 2018). The earliest presence of modern humans in SEA is dated to ∼50 thousand years ago (kya) (Higham 2013; Bae et al. 2017), followed by Paleolithic migration to East Asia ∼30 kya, inferred from genetic data (Yan et al. 2014; Hallast et al. 2015). There was also an expansion of Neolithic farmers and Bronze Age migrations from southern China to MSEA, which contributed to the present-day gene pool of modern MSEA people, e.g. Thais and Laotians (Higham 2014; Higham 2017; Lipson et al. 2018; McColl et al. 2018). Additional migrations during the historical period from neighboring countries (Penth 2000; Schliesinger 2000) have additionally enhanced ethnolinguistic diversity.

**Figure 1.**
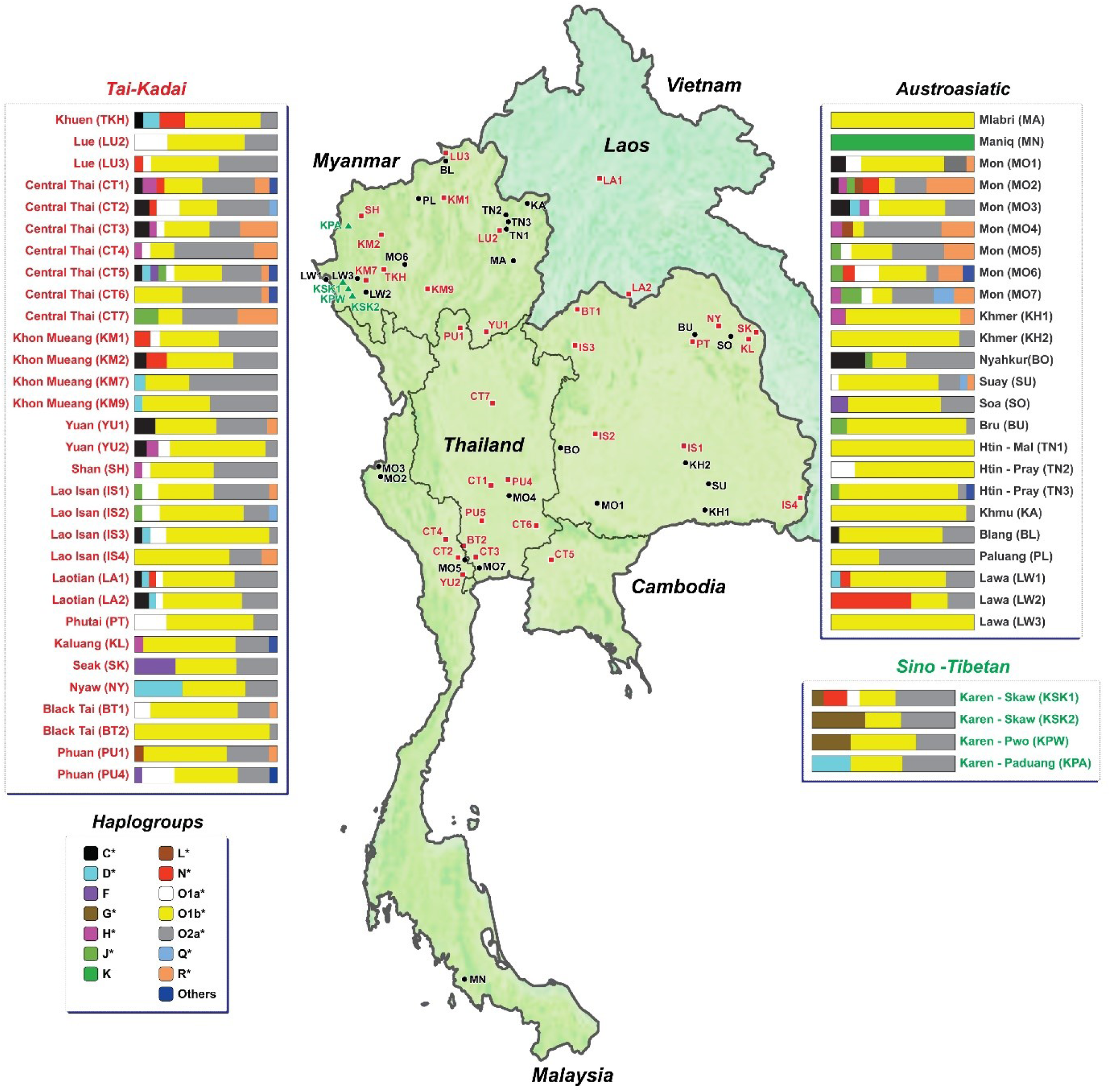
Map showing sample locations and haplogroup distributions.

The census size for Thailand was ∼68.41 million in 2017, and for Laos was ∼6.76 million in 2016 (Simons and Fennig 2018). There are five linguistic families distributed in these two countries. While the Tai-Kadai (TK) language is widely spread in southern China and MSEA, it is concentrated in present-day Thailand and Laos as it is a major language spoken by Thais (90.5%) and Laotians (67.7%). Austroasiatic (AA) speakers are next most frequent, accounting for 4.0% in Thailand and 24.4% in Laos. In addition, this area is also inhabited by historical migrants who speak Sino-Tibetan (ST), Hmong-Mien (HM), and Austronesian (AN) languages (frequencies of 3.2%, 0.3% and 2% respectively in Thailand; 3.1%, 4.8% and 0% in Laos) (Simons and Fennig 2018).

It is generally thought that AA languages were brought to the Thai/Lao region by Neolithic farmers from southern China, while TK languages were brought by a later, Bronze Age migration, also from southern China (Bellwood 2018). The Neolithic expansion was ∼2-3 kya before the expansion of TK languages; thus, the AA people were thought to be present before the TK expansion. The TK migration during the Bronze Age could have occurred via either demic diffusion (an expansion of TK people that brought both their genes and their language) or cultural diffusion (a language spread with minor movement of people). A genetic study on the origin of TK people supports a southern Chinese origin (Sun et al. 2013), while our previous studies of mtDNA genome sequences supports demic diffusion as the best explanation for the origin of the present-day Thai/Lao TK groups, although there is a strong signal of admixture between TK and AA groups in central Thailand (Kutanan et al. 2017; Kutanan et al. 2018b).

The male-specific portions of the Y chromosome (MSY) are paternally-inherited and exhibit lineages specific to populations/geographic regions, making the MSY an informative tool for reconstructing paternal genetic history and demographic change (Barbieri et al., 2014; Yan et al., 2014). However, to date there have been few MSY studies of MSEA and almost all of them employed short tandem repeats (Y-STRs) (Cai et al. 2011; Kutanan et al. 2011; Brunelli et al. 2018) which complicates comparison with mtDNA sequences due to their different mutational mechanism. In addition, those previous studies have also defined haplogroups by genotyping assays, which are thus biased in terms of the haplogroups detected and cannot uncover new sublineages.

We have previously carried out comprehensive studies of the maternal genetic history of the Thai/Lao region, based on 1,823 complete mtDNA genome sequences (Kutanan et al. 2017; Kutanan et al. 2018a; Kutanan et al. 2018b). In order to investigate the paternal genetic variation and demographic history, here, we investigate ∼2.3 mB of MSY sequence in a subset of the above individuals, comprising 928 sequences from 59 populations. We compare and contrast the MSY and mtDNA results, and we also use demographic modeling to address the role of demic vs. cultural diffusion vs. admixture in the origins of the major TK groups in each Thai/Lao region. Our MSY sequencing results provide new insights into the paternal genetic history of MSEA, and indicated contrasting paternal and maternal histories in this region.

## Results

We generated 914 sequences of ∼2.3 mB of the MSY, which combined with 14 published sequences brings the total to 928 MSY sequences belonging to 59 populations from Thailand and Laos (Figure 1; Table S5). There are 816 haplotypes defined by 8160 polymorphic sites, with mean coverages ranging from 4X to 109X (overall average coverage = 23X). Among the 928 MSY sequences, there are 92 specific haplogroups, belonging mostly to two main MSY lineages (O1b* and O2a*), that contribute substantially to the paternal genetic makeup of Thailand and Laos. There are several subclades of O1b*; the most frequent (50.54%) is O1b1a1a* or O-M95*, which occurs in almost half of the AA groups with a very high frequency (>70%), i.e. KH1-KH2, KA, BU, BL, SU, TN1-TN3, MA and LW3 (Figure 1: Table S1). The Correspondence Analysis (CA) (based on haplogroup frequencies) also supports the divergence of these AA speaking groups in agreement with the other results mentioned later, with many O1b* sublineages, e.g. O1b1a1a1b1a (O-B426) and O1b1a1a1a1a (O-F2758) (Figure S1). O2a* or O-M324* is the second most frequent haplogroup (25.86%) and has a relatively high frequency (>40%) in some AA and TK groups, and all ST speaking Karen. Additional minor non-SEA specific haplogroups were also observed, e.g. haplogroup N, found in the Lawa groups, and haplogroups R*, H*, and J*, which support associations between Indian and the Mon, and genetic connections between Mon and TK groups (Figure 1 and Figure S1). Further details on haplogroup distribution are provided in Table S1 and Supplementary Text.

### Genetic diversity and structure

Generally, the AA populations show lower genetic diversity values than the TK and ST groups for the MSY, in agreement with the mtDNA results (Figure 2) (Mann–Whitney *U* tests between AA and TK for MSY: *h*: *Z* = 3.37, *P* < 0.01, MPD: *Z* = 2.40, *P* < 0.05, haplogroup diversity: Z = 3.74, *P* < 0.01 and for mtDNA: *h*: *Z* = 4.33, *P* < 0.01, MPD: *Z* = 1.47, *P* > 0.05, haplogroup diversity: *Z* = 4.37, *P* < 0.01). After the Maniq (MN), who have no MSY variation, and the Mlabri (MA), who have no mtDNA variation, the Htin (TN1), Lawa (LW3) and Bru (BU) show very low diversity values of MSY whereas the Htin (TN1-TN3), Khmer (KH2) and Seak (SK) show low mtDNA diversity (Figure 2). In contrast to the other AA groups, the Mon (MO1-MO7) show higher levels of both MSY and mtDNA diversity than other AA groups (Mann–Whitney *U* tests between AA and Mon for MSY: *h*: *Z* = −3.33, *P* < 0.01, MPD: *Z* = −3.30, *P* < 0.01, haplogroup diversity: *Z* = −3.75, *P* < 0.01 and for mtDNA: *h*: *Z* = −1.94, *P* > 0.05, MPD: *Z* = −2.03, *P* < 0.05, haplogroup diversity: *Z* = −2.79, *P* < 0.01). LW3 showed very low MSY haplogroup diversity and MPD values (Figure 2C), and a significantly low Tajima’s D value (Figure 2D), suggesting recent paternal expansion in this group, but the converse trend (rather high diversity) for mtDNA. Interestingly, a significantly negative Tajima’s D value was observed more frequently in the TK than the AA groups for both the MSY and mtDNA (MSY, *P* < 0.05: 10/31 for TK vs. 6/24 for AA; mtDNA, *P* < 0.05: 20/31 for TK vs. 5/24 for AA) (Figure 2D), suggesting a stronger signal of recent population expansion in TK groups; no significant Tajima’s D values were observed in any of the ST-speaking Karen groups. The Nyahkur (BO), who speak a Mon language, show the highest MPD value for the MSY, which might indicate paternal gene flow with other populations; this is supported by the BO having the highest number of shared MSY haplotypes (3 haplotypes) with other populations (Figure 3A). MO3 and MO4 have shared MSY haplotypes with the TK speaking groups (CT2, CT6 and YU1), reflecting their genetic connection. In the mtDNA, apart from the AA-speaking Palaung (PL), the Mon (MO2, MO3 and MO7) also share haplotypes with the central Thai (CT3 and CT6) and Shan (SH) (Figure 3A).

**Figure 2.**
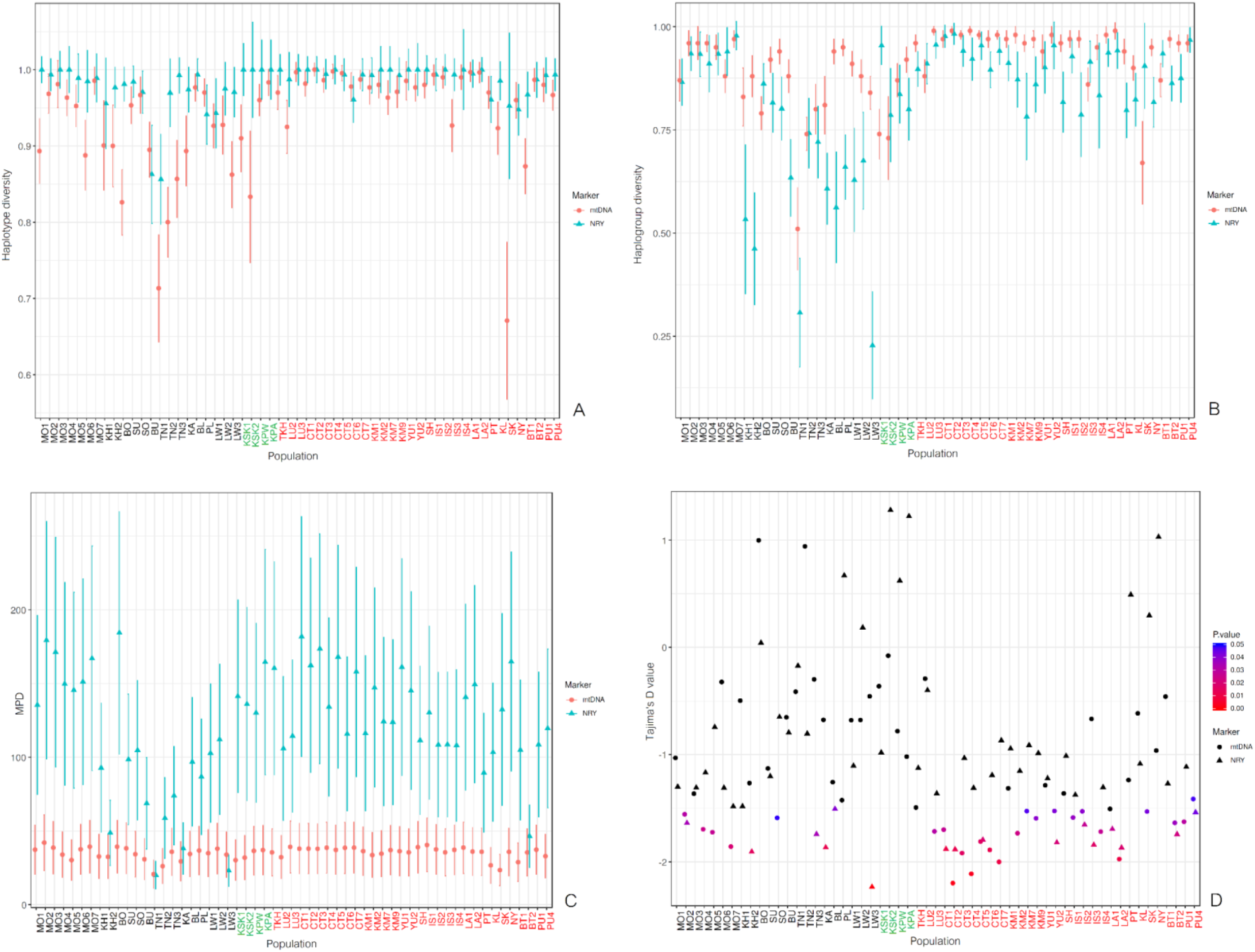
Genetic diversity values of MSY and mtDNA in the studied populations, excluding the Maniq (MN) and Mlabri (MA): haplotype diversity (A), haplogroup diversity (B), mean number of pairwise difference (MPD) (C), and Tajima D’s values (D). More information and all genetic diversity values are provided in Table S5.

**Figure 3.**
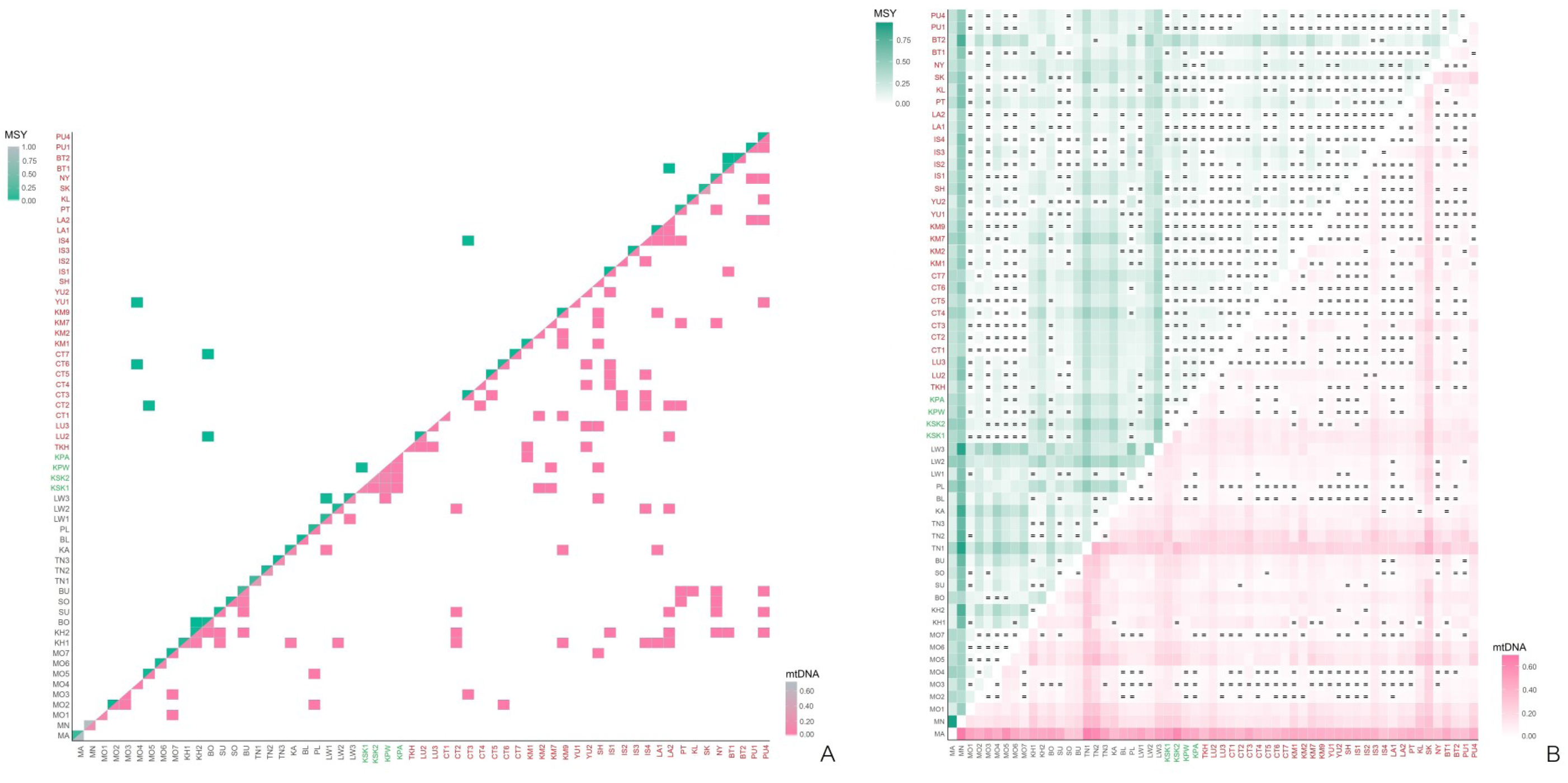
Relative shared haplotypes (A) and heat plot of *Φ*_*st*_ (B) between studied populations for the MSY and for mtDNA.

The Analysis of Molecular Variance (AMOVA) indicates that the variation among groups accounts for 11.20% of the total MSY genetic variance (Table 1). There is greater genetic heterogeneity among the AA groups (20.01%, *P* < 0.01 and 18.49%, *P* < 0.01 without MN, the hunter-gatherer group from southern Thailand) than among the TK (4.48%, *P* < 0.01) and ST-speaking Karen groups (2.29%, *P* > 0.01). For the AA groups with more than one population sampled, the greatest among-group variation by far was among the three Lawa populations (34.43%, *P* < 0.01), while the seven Mon populations showed very low (albeit still significant) among-group variation (3.92%, *P* < 0.01) (Figure S2). Very low among-group variation was also observed for the central Thai groups from central Thailand (1.47% *P* > 0.01), Khon Mueang groups from northern Thailand (−1.83%, *P* > 0.01), and Lao Isan groups from northeastern Thailand (1.84%, *P* > 0.01), indicating overall genetic homogeneity among these major TK speaking groups. In agreement with the MSY, larger mtDNA variation is observed in the AA groups (14.03%, *P* < 0.01) than the ST (6.51%, *P* < 0.01) and TK groups (4.33%, *P* < 0.01), but interestingly the largest among-group variation is not among the Lawa (7.78%, *P* < 0.01) but rather among the Htin populations (25.71%, *P* < 0.01). In contrast to the MSY, each of the TK groups with more than one population sampled showed significant among-group differences for mtDNA, especially the Khon Mueang (4.20%, *P* < 0.01) (Figure S2). In sum, we observed different patterns of MSY vs. mtDNA for the different language groups. The among-population variation within linguistic groups is larger for the MSY (20.01%, *P* < 0.01) than for mtDNA (14.03%, *P* < 0.01) for AA groups, but about the same for TK groups (4.48%, *P* < 0.01 for MSY and 4.33%, *P* < 0.01 for mtDNA), and the ST groups have larger among-population variation for mtDNA (6.51%, *P* < 0.01) than for the MSY (2.29%, *P* < 0.01) (Table 1; Figure S2). Thus, there are different patterns of MSY vs. mtDNA differentiation for these three language families.

**Table 1.**
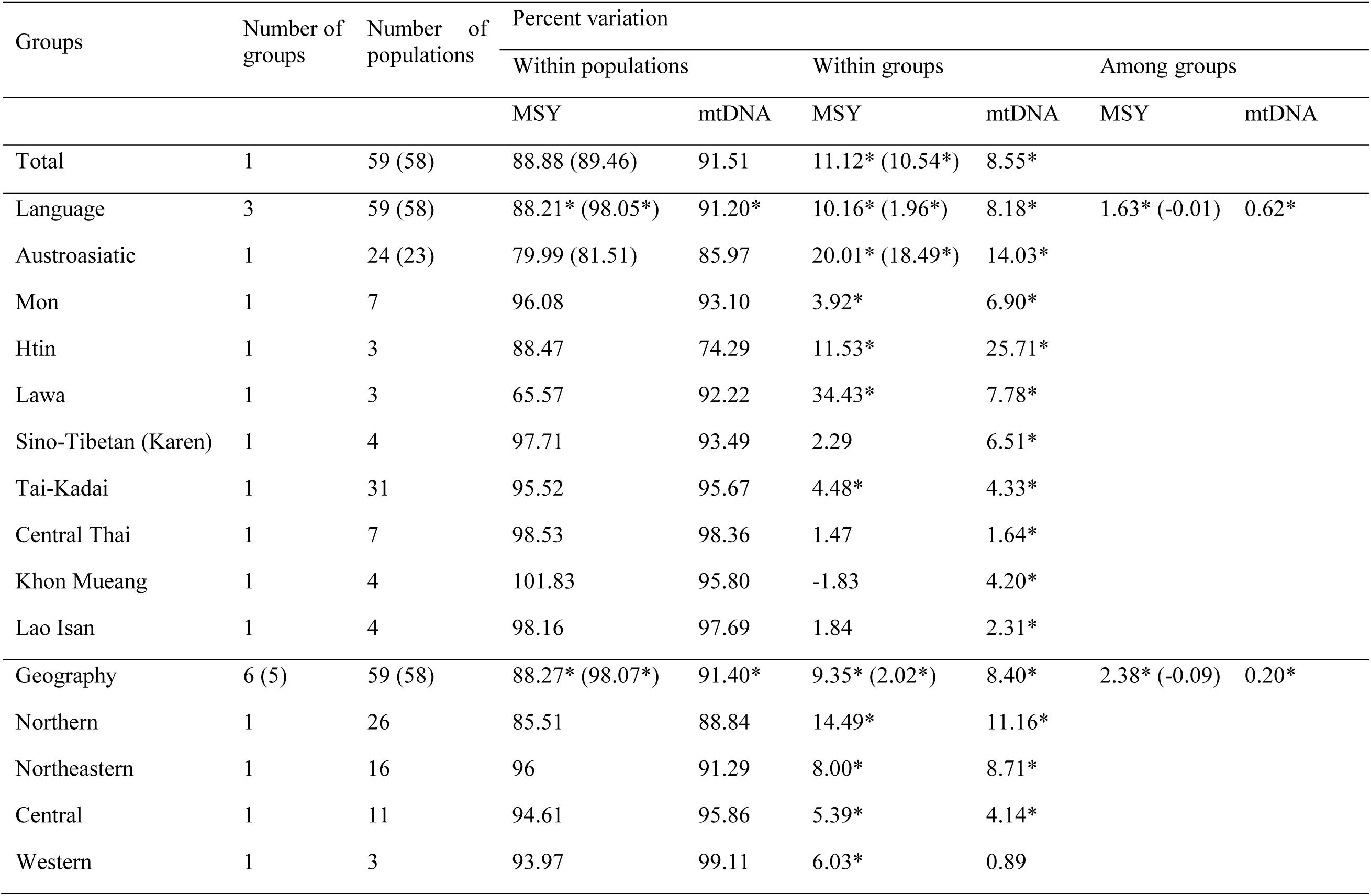
AMOVA results. The numbers in parentheses show the percent variation of MSY by excluding the Maniq (MN) and asterisks indicate significant level (*P* < 0.01).

| Groups | Number of groups | Number of populations | Percent variation |  |  |  |  |  |
| --- | --- | --- | --- | --- | --- | --- | --- | --- |
|  |  |  | Within populations |  | Within groups |  | Among groups |  |
|  |  |  | MSY | mtDNA | MSY | mtDNA | MSY | mtDNA |
| Total | 1 | 59 (58) | 88.88 (89.46) | 91.51 | 11.12* (10.54*) | 8.55* |  |  |
| Language | 3 | 59 (58) | 88.21* (98.05*) | 91.20* | 10.16* (1.96*) | 8.18* | 1.63* (-0.01) | 0.62* |
| Austroasiatic | 1 | 24 (23) | 79.99 (81.51) | 85.97 | 20.01* (18.49*) | 14.03* |  |  |
| Mon | 1 | 7 | 96.08 | 93.10 | 3.92* | 6.90* |  |  |
| Htin | 1 | 3 | 88.47 | 74.29 | 11.53* | 25.71* |  |  |
| Lawa | 1 | 3 | 65.57 | 92.22 | 34.43* | 7.78* |  |  |
| Sino-Tibetan (Karen) | 1 | 4 | 97.71 | 93.49 | 2.29 | 6.51* |  |  |
| Tai-Kadai | 1 | 31 | 95.52 | 95.67 | 4.48* | 4.33* |  |  |
| Central Thai | 1 | 7 | 98.53 | 98.36 | 1.47 | 1.64* |  |  |
| Khon Mueang | 1 | 4 | 101.83 | 95.80 | -1.83 | 4.20* |  |  |
| Lao Isan | 1 | 4 | 98.16 | 97.69 | 1.84 | 2.31* |  |  |
| Geography | 6 (5) | 59 (58) | 88.27* (98.07*) | 91.40* | 9.35* (2.02*) | 8.40* | 2.38* (-0.09) | 0.20* |
| Northern | 1 | 26 | 85.51 | 88.84 | 14.49* | 11.16* |  |  |
| Northeastern | 1 | 16 | 96 | 91.29 | 8.00* | 8.71* |  |  |
| Central | 1 | 11 | 94.61 | 95.86 | 5.39* | 4.14* |  |  |
| Western | 1 | 3 | 93.97 | 99.11 | 6.03* | 0.89 |  |  |

Although there is more variation among groups defined by geographic location (2.38%, *P* < 0.01) than by language family (1.63%, *P* < 0.01) (Table 1) there is much more MSY variation among populations within the same group than among groups defined either by geographic or linguistic criteria. Moreover, when the divergent MN population of hunter-gatherers from southern Thailand is removed from the analysis, then the among-group component is no longer significant for either geographic location or language family (−0.09%, *P* > 0.01 for geography; −0.01%, *P* > 0.01 for language), and the total variation among populations reduces to 10.54%. Thus, neither geography nor language family is a good predictor of the MSY genetic structure of Thai/Lao populations.

There are significant correlations between matrices of MSY genetic and geographic distance, estimated by Mantel tests, for all three types of geographic distances, i.e. great circle distance (*r* = 0.3381, *P* < 0.01), resistance distance (*r* = 0.5418, *P* < 0.01) and least-cost path distance (*r* = 0.3912, *P* < 0.01). However, the correlations are no longer significant when the MN group is removed from the analysis: great circle distance (*r* = 0.0125, *P* > 0.05), resistance distance (*r* = −0.0446, *P* > 0.05) and least-cost path distance (*r* = 0.0139, *P* > 0.05). In contrary, no significance was detected (*P* > 0.05) between matrices of mtDNA genetic distance and geographic distances with and without MN (great circle distance: *r* = 0.0776 and *r* = −0.0323), resistance distance (*r* =0.1433 and *r* = −0.1105) and least-cost path distance (*r* = 0.0997 and *r* = −0.0253).

To identify and describe population clustering based on multivariate analysis, Discriminant Analysis of Principal Components (DAPC) was carried out. This analysis attempts to maximize among-groups genetic differentiation and minimize within-group genetic variation; the results showed considerable overlap among groups defined by either language family or geographic location in both MSY and mtDNA (Figure S3). In addition, the groupings by population and ethnicity of MSY data revealed the largest discrimination to be among some AA-speaking groups, i.e. all Lawa groups (LW1-LW3), Htin (TN1) and Blang (BL) whereas all Htin groups (TN1, TN2 and TN3), Mlabri (MA), TK-speaking Seak (SK) and ST-speaking Karen (KSK1, KSK2 and KPW) are differentiated from the others for mtDNA, emphasizing contrasting genetic pattern between MSY and mtDNA for Htin, Mlabri, Lawa, Blang, Seak and Karen.

In sum, all results indicate lower genetic diversity of the AA groups than the TK and ST groups, except the Mon and Nyahkur, who exhibit high genetic diversity. The AA groups also show greater genetic heterogeneity than the TK and ST groups.

### Post-marital residence and genetic diversity

Although the influence of post-marital residence (patrilocal vs. matrilocal) has previously been studied in northern Thai hill tribes, these studies compared genetic variation between partial mtDNA sequences (hypervariable regions of the control region) and Y-STR loci (Oota et al. 2001; Besaggio et al. 20007). Here, we report the first comparison of mtDNA and MSY variation based on comparable sequence data. We studied four hill tribes (Karen, Htin, Lawa and Khmu) and the Palaung, another minority group in the mountainous area of northern Thailand but not officially recognized as a hill tribe. The Khmu (KA), Lawa (LW1, LW2 and LW3) and Palaung (PL) groups practice patrilocality (i.e., the wife moves to the residence of her husband after marriage), whereas the Htin (TN1, TN2 and TN3) are matrilocal, as are the ST-speaking Karen (KSK1, KSK2, KPA and KPW). If residence pattern is influencing genetic variation, then lower within-population genetic diversity coupled with greater genetic heterogeneity among populations is expected for patrilocal groups than for matrilocal groups for the MSY, while the opposite pattern is expected for mtDNA (Oota et al. 2001). Mean values of *h*, MPD and haplogroup diversity in MSY are higher in matrilocal than patrilocal groups, not significantly different for *h* and MPD (Mann–Whitney *U* tests: *h*: *Z* = 1.4616, *P* > 0.05; MPD: *Z* = 0.9744, *P* > 0.05) but significantly different for haplogroup diversity (Mann–Whitney *U* tests: Z = 2.1112, *P* < 0.05) (Figure S4). For mtDNA, non-significant higher genetic diversity values for patrilocal than matrilocal groups are observed (Mann–Whitney *U* tests: *h*: *Z* = −0.9744, *P* > 0.05; MPD: *Z* = −0.8120, *P* > 0.05; haplogroup diversity: z = −1.864, *P* > 0.05) (Figure S4). Notably, TN1 and LW3 exhibit very low with-in population diversity for the MSY, e.g. MPD = 20.07 and 23.07, compared to the average MPD (121.11), whereas TN1 and TN2 (20.69 and 26.14) show lower MPD than average (35.09) for mtDNA. For genetic differences between-populations, the patrilocal Khmu, Lawa and Palaung have significantly higher genetic differentiation for the MSY than for mtDNA (average *Φ*_*st*_ = 0.3109 for MSY and 0.0774 for mtDNA) (Mann–Whitney *U* tests: *Z* = 3.5907, *P* < 0.01) whereas the matrilocal groups (Htin and Karen) also show higher average *Φ*_*st*_ for MSY (0.1859) than for mtDNA (0.1553), but these are not significantly different (Mann–Whitney *U* tests: *Z* = 0.3270, *P* > 0.05). Contrasting genetic differences for the MSY vs. mtDNA of Lawa, Htin and Karen are clearly seen in the MDS and DAPC plots (Figure 4A and 4B; Figure S3). Much stronger contrasting between-group variation is seen in the AMOVA results (Lawa: 34.43% for MSY and 7.78% for mtDNA; Htin: 11.53% for MSY and 25.71% for mtDNA; Karen: 2.29% for MSY and 6.51% for mtDNA (Table 1; Figure S2).

**Figure 4.**
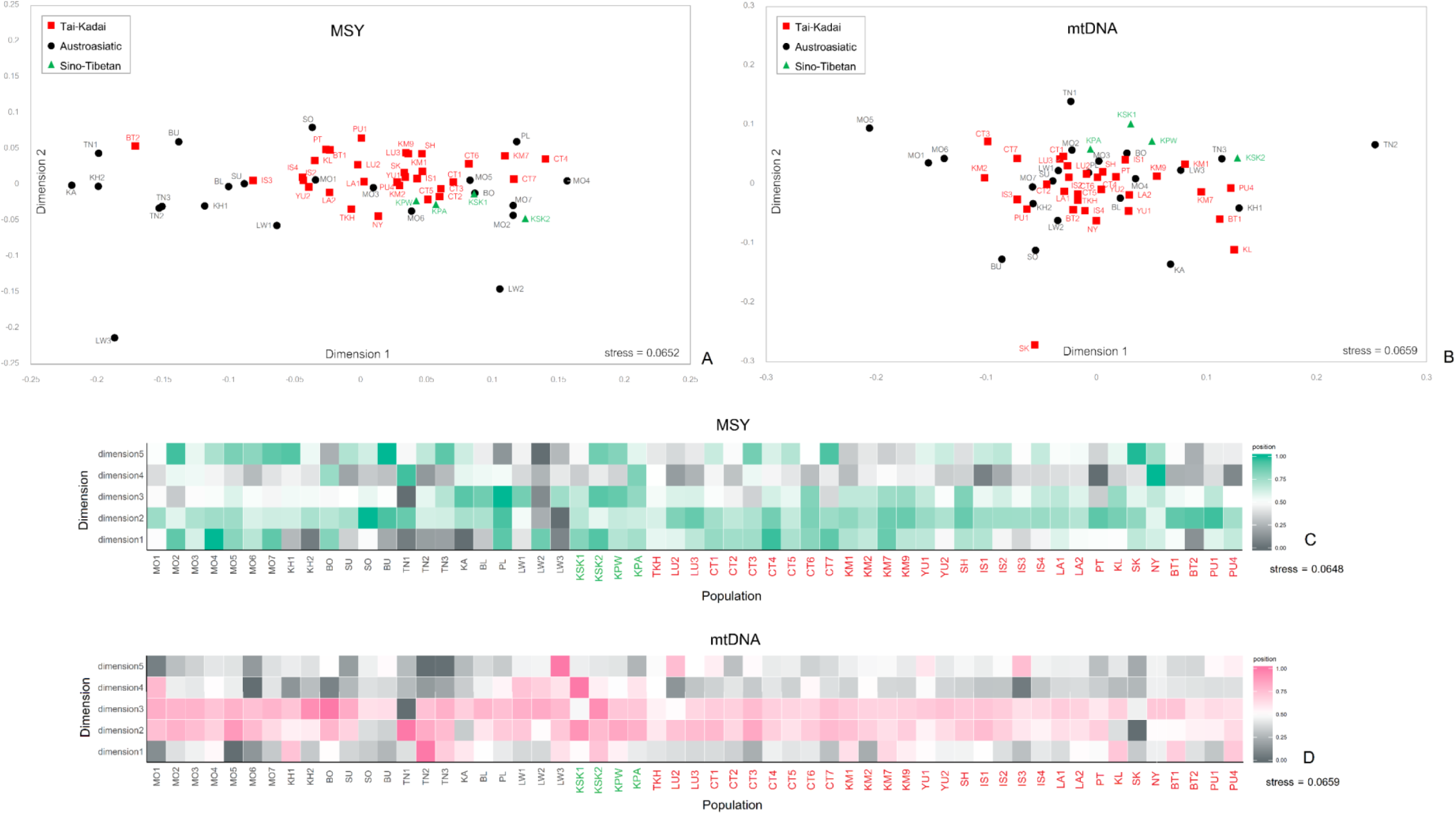
The two-dimensional MDS plot and five-dimensional MDS heat plot based on the *Φ*_*st*_ distance matrix for 57 populations (after removal of Maniq and Mlabri) of MSY (A and C) and mtDNA (B and D).

However, in general, the AA-speaking groups, whether identified as hill-tribes or as other minorities, are patrilocal groups. The AMOVA result indicates that the variation among AA populations is higher in MSY (20.01%) than mtDNA (14.03%), in accordance with expectations if residence pattern is influencing genetic variation. Conversely, the TK populations, where neither patrilocal nor matrilocal residence is preferred, exhibit similar among-population variances for the MSY (4.48%) and mtDNA (4.33%) (Table 1; Figure S2). Overall, there does seem to be some impact of post-marital residence on the patterns of genetic diversity.

### Genetic relatedness among populations

The genetic distance and MDS analyses based on MSY and mtDNA indicate that the MN and MA are highly diverged from the other populations for the MSY and mtDNA, respectively (Figure 3B and S5). The MA and MN also show large differences from the other populations in the heat plots of *Φ*_*st*_ values (Figure S5). However, in general both MSY and mtDNA results show relatively larger genetic heterogeneity of the AA groups vs. genetic homogeneity of the TK and ST groups (Figure 3B). The Mantel test of *Φ*_*st*_ values showed a significant correlation between the MSY and mtDNA *Φ*_*st*_ matrices (*r* = 0.4506, *P* < 0.01). After excluding these MA and MN as outliers, the MDS for the MSY showed that almost all AA speaking groups are located along the edges of the plot, while most of the TK groups cluster in the center of the plot (Figure 4A), further supporting genetic heterogeneity of the AA and homogeneity of the TK populations. Interestingly, the SEA-specific O-M95* and O-M234* haplogroups (with several sublineages) differentiate the studied populations into at least two main paternal sources, and the frequencies of these two haplogroups correspond to the major differentiation in the MDS plot (Figure 4A). O-M95* is at high frequency in the populations on the left of the plot and gradually decreases to very low frequency in the populations on the right side in the first dimension, whereas the O-M324* frequency runs opposite to the O-M95* cline: O-M324* is at higher frequency in populations located on the right of the plot and decreases in frequency toward the left side (Figure 4A). The MDS plot and heat plot of MSY also indicates some Mon groups (MO1, MO3, MO5 and MO6) are close to the cluster of TK groups in the center of the plot (Figure 4A and 4C), indicating a close genetic relationship. In addition, non-SEA haplogroups lineages, e.g. R*, H*, and J*, provide more support for genetic connections between Mon and Central Thais.

For the MDS based on mtDNA (Figure 4B), the Mon generally showed genetic affinities with the TK groups in the center of the plot, with the exception of MO1, MO5 and MO6, which differ from the other Mon groups, as can be also seen in the MDS plot and heat plot (Figure 4B and 4D). Overall, we observe more genetic heterogeneity of the AA groups than the other linguistic groups and there are contrasting patterns of genetic relationships for the MSY vs. mtDNA.

### Genetic relatedness between Thai/Lao and other Asian populations

The MDS based on the MSY *Φ*_*st*_ matrix of 73 populations from across Asia revealed that, in general, population clustering largely reflects linguistic affiliation (Figure 5), with some exceptions. In the first and second dimension, the AA populations are the most diversified, with the PL and MN appearing as outliers. There is one cluster of AA populations on the left, which also includes one TK group (BT2); the other AA populations are scattered along the main axis of the plot. Some Mon groups (MO2, MO4 and MO7) are relatively close to Indian and ISEA populations, indicating potential connections. Two central Thai groups (CT4 and CT7) are also relatively close to the Indian populations. The ST populations (Karen, Han Chinese and Burmese) are rather close. The ISEA and Papuan populations are in closer proximity to South Asian populations (Indian, Bengali and Punjabi). Generally, the haplogroup profile indicates genetic affinities between the Mon and South/Central Asian groups, which is consistent with the MDS plots (Figure 5) and results from previous mtDNA haplogroup analyses (Kutanan et al. 2017; Kutanan et al. 2018b).

**Figure 5.**
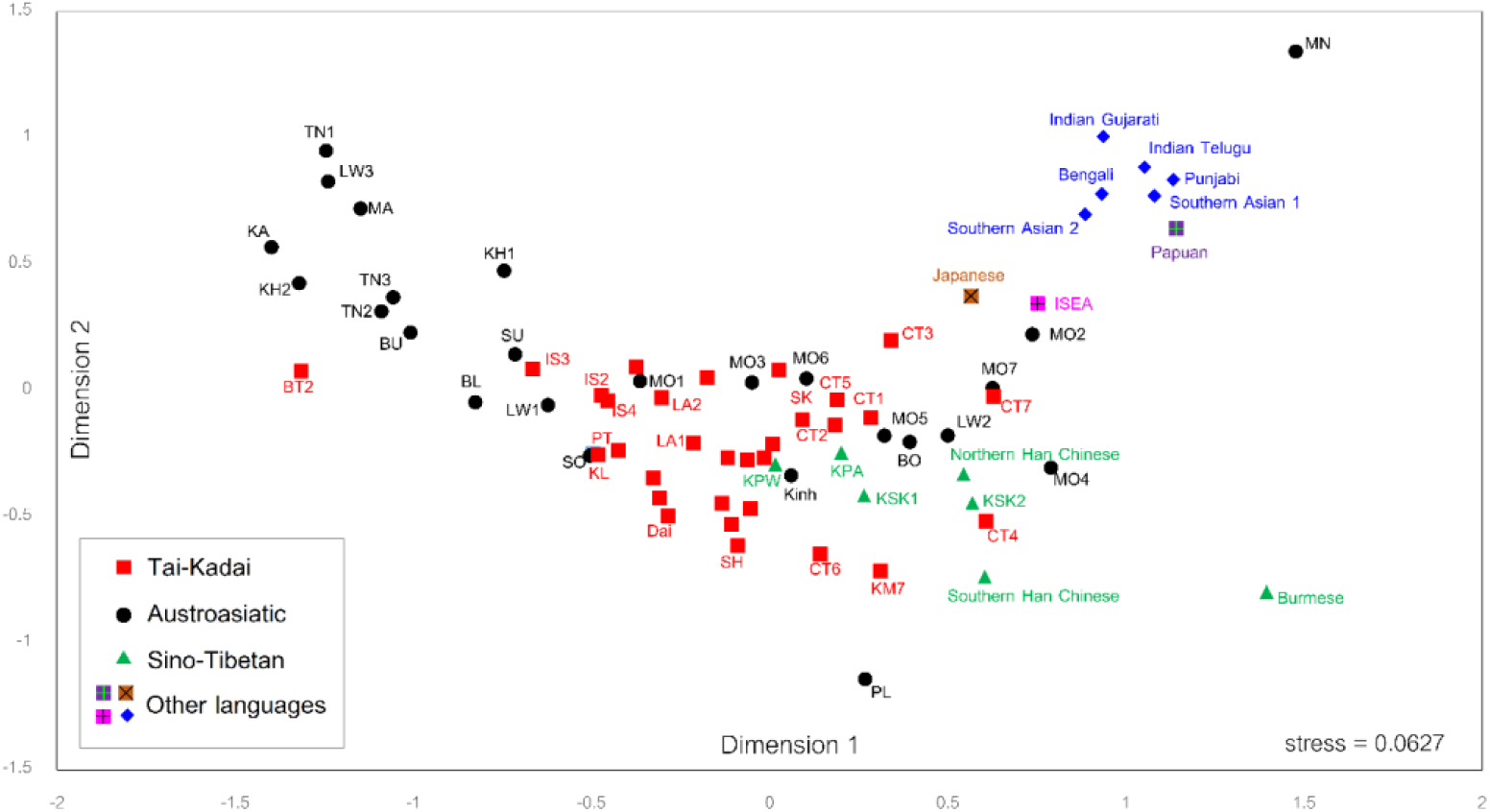
The two-dimensional MDS plot based on the MSY *Φ*_*st*_ distance matrix for 73 populations. Population details are listed in Figure 1 and Tables S5 and S7.

### The expansion of male lineages

The Bayesian Skyline Plots (BSP) of effective population size change (*N*_e_) over time in each group reveal overall 5 different trends (Figure 6). The most common trend, found in Mon, Khmer, Htin, Central Thai and Black Tai, showed *N*_e_ increasing gradually or remaining constant during 40-60 kya until a decline ∼5-7 kya, followed by rapid growth ∼5 kya and then a decrease ∼2.0-2.5 kya. The other trends differ from the first trend as follows: no population reduction ∼2.0-2.5 kya but population size either increases (Khon Mueang and Yuan) or remains stable (Lao Isan and Laotian); the Lue and Phuan show two increases in *N*_e,_ at about ∼5 kya and ∼10 kya; the Lawa show a stable population size since ∼30 kya and then a decline during the last 2 kya with a sudden increase ∼1 kya; and the Karen differ only slightly from the common trend with a population increase ∼1 kya.

**Figure 6.**
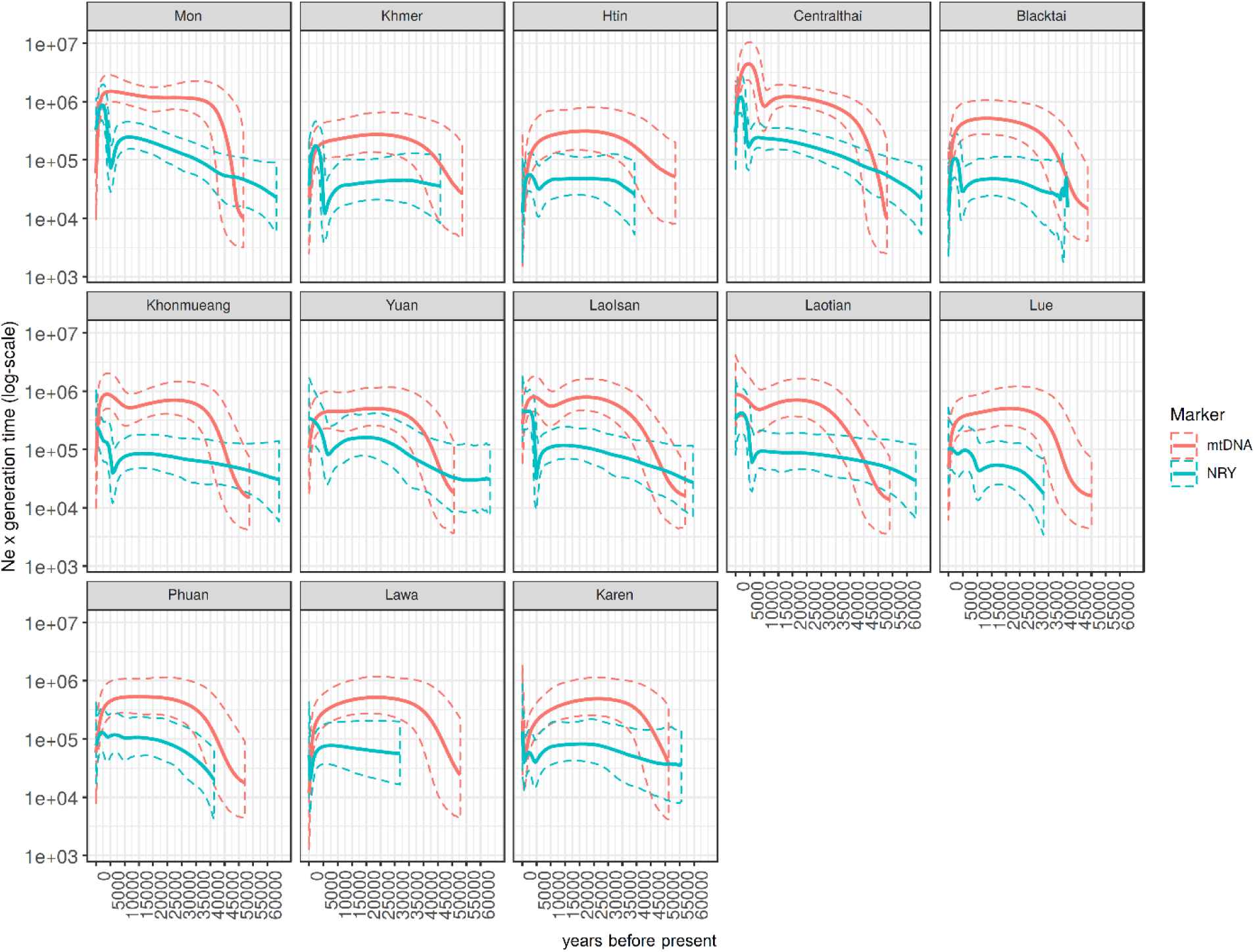
The BSP plots based on the MSY and mtDNA of 13 ethnicities from Thailand and Laos; Mon, Khmer, Htin, Central Thai, Black Tai, Khon Mueang, Yuan, Lao Isan, Laotian, Lue, Phuan, Lawa, Karen. Solid lines are the median estimated effective population size (y-axis) through time from the present in years (x-axis). The 95% highest posterior density limits are indicated by dotted lines.

By contrast, the BSP based on mtDNA sequences for each ethnicity show three common trends (Figure 6). The first trend is an increase in *N*_e_ during 40-50 kya, followed by stability and then decrease ∼2 kya, which was observed in Mon, Htin, Lawa, Khmer, Yuan, Phuan and Lue. The second pattern, shown by the Khon Mueang, is an increase in *N*_e_ ∼40-50 kya, followed by stability and then increase again ∼10 kya, followed by a decline ∼2 kya. The Central Thai, Lao Isan and Laotian show the third trend, in which population increases occur ∼40-50 and ∼10 kya. In general, the BSP plot by ethnicity indicated lower effective population sizes for the MSY than for mtDNA (Figure 6).

We also plotted the BSP of several Asian populations from published MSY data (Karmin et al. 2015; Poznik et al. 2016) (Figure 7). Almost all of the MSEA and East Asian populations, i.e. Kinh, northern Han, southern Han and Japanese show a pronounced increase of the MSY *N*_e_ during ∼4-6 kya, except the Xishuangbanna Dai, in which there is an increase ∼2 kya. Around 5 kya, the Japanese show a decrease in *N*_e_ before a sudden increase, suggesting a bottleneck prior to demographic expansion. Interestingly, the ISEA population shows a large increase in *N*_e_ ∼35-40 kya and a smaller increase ∼2.5-3 kya. The South Asian populations, i.e. Bengali, Punjabi and Indian, also show two pulses of population increase at about the same times. The Punjabi also show an additional small increase in *N*_e_ change during ∼12 kya.

**Figure 7.**
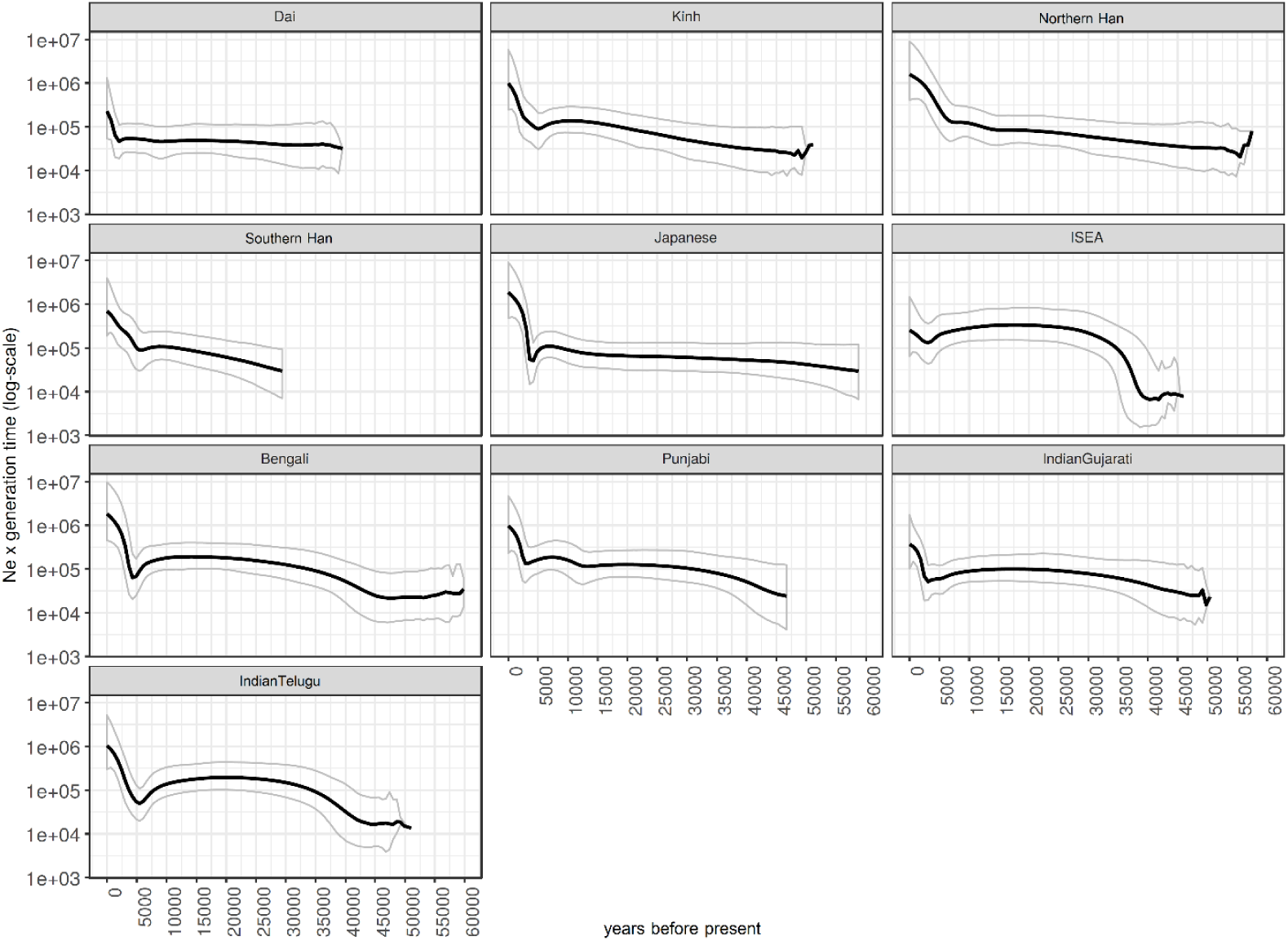
The BSP plots of Asian populations. Solid lines are the median estimated paternal effective population size (y-axis) through time from the present in years (x-axis). The 95% highest posterior density limits are indicated by dotted lines.

The BSP by each major MSY haplogroup show four pulses of paternal *N*_e_ increases, at ∼9-11 kya, ∼5 kya, ∼2.0-2.5 kya and ∼1.0 kya (Figure 8), in agreement with the plot by ethnicity. The early Holocene *N*_e_ increment is obviously noticed in O2a1c* and O2a2a*, whereas the *N*_e_ growth ∼5 kya is observed in O1b1a1a1b* and R*. Haplogroup O1a*, C* and D* show expansions in *N*_e_ ∼2.0-2.5 kya and haplogroup N* shows a recent expansion ∼1.0 kya. In addition, there are two expansion times for O1b1a1a1a* and O2a2b* (∼5 and ∼2 kya).

**Figure 8.**
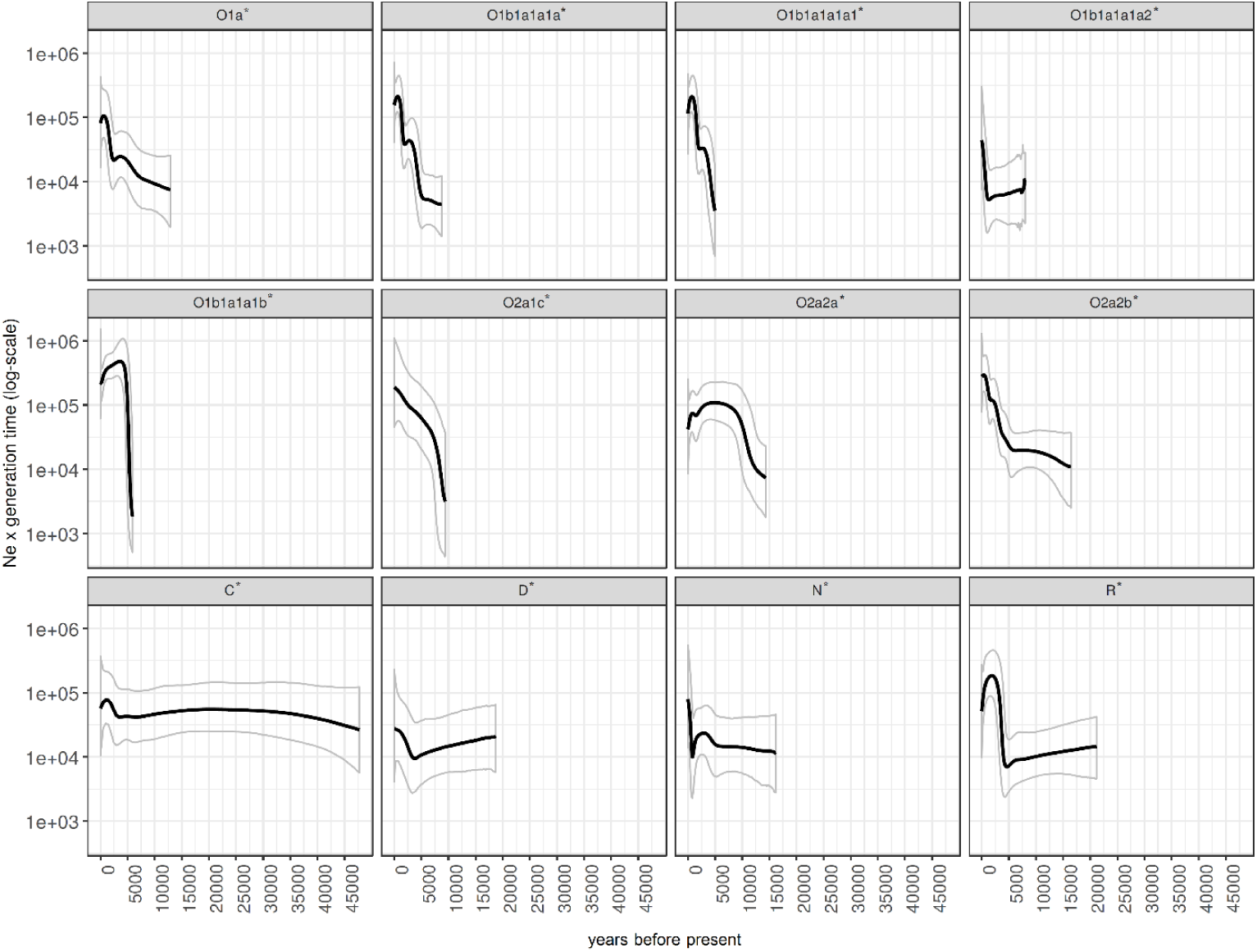
The BSP plots for each major haplogroup. Solid lines are the median estimated paternal effective population size (y-axis) through time from the present in years (x-axis). The 95% highest posterior density limits are indicated by dotted lines.

### Demographic models

Previously, we used mtDNA genome sequences and demographic modeling to test different hypotheses about the origins of TK groups. Specifically, we tested whether different TK groups were primarily related to local AA groups (reflecting cultural diffusion, i.e. an AA group switching to a TK language), to a TK group from southern China (reflecting demic diffusion, i.e. spread of TK languages via migration from southern China), or were related to both (reflecting admixture between an incoming TK group from southern China and a local AA group). We found that the Khon Mueang (from northern Thailand), Lao Isan (from northeastern Thailand) and Laotian most likely originated via demic diffusion from southern China without substantial gene flow from AA groups (Kutanan et al. 2017). However, for the central Thai, the most likely scenario was demic diffusion with a very low level of gene flow between central Thai and Mon groups (Kutanan et al. 2018b). Here we use the same approach to test three demographic scenarios concerning the paternal origins of these major Thai groups (Figure S6).

For the Khon Mueang (KM) people (Test 1), the highest posterior probability (0.80) and rather highly selected classification trees (0.58) were found for the demic diffusion model (Table S2). By contrast, the cultural diffusion model is the most likely scenario for the Lao and central Thai groups. Both the combined Laotian (LA) and Lao Isan (IS) datasets (Test 2) and the separate LA dataset (Test 3) weakly support the cultural diffusion model (for Test 2; posterior probability = 0.56 and selected classification tree = 0.37 and for Test 3; posterior probability = 0.56 and selected classification tree = 0.39). The IS dataset (Test 4) supports cultural diffusion (with the present-day IS groups descended from local Khmer (KH) with the highest posterior probability (0.71) and classification trees selected slightly more often than for the other models (0.49). For Test 5 (the Central Thai (CT) dataset), the cultural diffusion model had the highest posterior probability at 0.58 and was selected slightly more often among the classification trees (0.50) than the other models. However, a Principal Component Analysis (PCA) plot shows that based on the first two PCs the observed data fall within the distributions simulated under the three models in only Test 4, whereas the other datasets fall within the simulated distributions for PCs 3 and 4, suggesting that there is low efficiency to reconstruct the variability of the observed data (Figure S7). The parameter estimation for the best performing models in all five tests was able to obtain point estimates for each of the simulated effective population sizes. However, the posterior distributions were generally flat (Table S3: Figure S8). We also calculated the MSY *Φ*_*st*_ and corrected pairwise difference among groups of populations used in ABC tests to estimate their genetic relationships (Table S4). The KM are closer to the Dai than the local AA group (Test 1), the ethnic Lao and Laotian showed similar genetic differences to both Dai and AA groups (Test 2 and Test 3), whereas the CT groups (Test 5) have closer genetic relationships to the local AA group than to Dai. In contrast, mtDNA *Φ*_*st*_ and corrected pairwise difference revealed that the KM and ethnic Lao are closer to the Dai than local AA while the CT exhibited somewhat similar genetic distances to both Dai and AA. Overall, the simulations based on MSY sequences, compared with previous mtDNA simulation together with tests of genetic difference by *Φ*_*st*_ and corrected pairwise differences, suggest different demographic histories for males and females in the region.

## Discussion

In order to gain more insights into MSEA genetic history, we here investigate the paternal genetic variation and structure by sequencing ∼2.3 mB of the MSY from representative ethnolinguistic groups from Thailand and Laos. In sum, most of the studied populations exhibit two major MSY haplogroups, O-M324* and O-M95* in different proportions, indicating two major paternal sources. O-M324* was widely spread in the TK groups, while O-M95* is predominant in the AA groups. However, some TK populations (BT2 and IS3) and some AA populations (PL, BO and MO4) exhibited the opposite pattern (Figure 1; Table S1). We also compared patterns of MSY variation with mtDNA in the same set of populations and found some similar results, e.g. overall lower genetic diversity and greater heterogeneity of AA groups than of TK and ST groups, large differences between the Mon and the other AA groups, and genetic connections between the Mon and central Thai (Figure 2-4). However, in many respects the patterns of MSY and mtDNA variation are different, suggesting contrasting paternal and maternal genetic histories.

### Factors influencing contrasting genetic variation in the hill tribes

Although the genetic variation of the studied populations appears to be influenced by both linguistic and geographic factors, when the very diverged Maniq group is removed, neither language nor geography impacts genetic variation, indicating that these two factors are not important in the broad view (Table 1). Other factors, i.e. cultural practices, admixture, and genetic drift, seem to be more influential. In the case of the hill tribes, post-marital residence and preservation of their identity are putatively influential factors. In Thailand, there are nine ethnic groups which are officially identified as hill tribes, i.e. the AA-speaking Lawa, Htin and Khmu, the HM-speaking Hmong and IuMien and the ST-speaking Karen, Lahu, Akha and Lisu. The Akha, Lisu, Hmong, IuMien, Lawa and Khmu practice patrilocality while the Lahu, Karen and Htin are matrilocal. If post-marital residence is influencing patterns of genetic variation, then the expectation is for larger between-group differences and smaller within-group diversity for patrilocal groups for the MSY, and the same trends for matrilocal groups for mtDNA (Oota et al. 2001). The first comparative study of mtDNA and MSY variation in patrilocal vs. matrilocal groups was carried out in northern Thai hill tribes, because they include both patrilocal and matrilocal groups within a small geographic scale, and found a strong impact of post-marital residence on the mtDNA and MSY variation (Oota et al. 2001). However, previous studies compared genetic variation between partial mtDNA sequences and Y-STRs (Oota et al. 2001; Besaggio et al. 20007); here we report the first comparison of mtDNA and MSY variation based on comparable sequence data.

We analyzed the Khmu, Palaung and Lawa groups, who practice patrilocality, whereas the Htin are matrilocal, similar to the ST-speaking Karen. The within-population genetic diversity values is in agreement with expectation, i.e. greater diversity of matrilocal than patrilocal groups for MSY and the opposite trend in mtDNA (Figure S4). Moreover, genetic differentiation between populations is correlated with post-marital residence. However, in many cases the differences between patrilocal and matrilocal groups are not significant, indicating that other factors are also having an effect. One factor in particular that could influence the within-population genetic diversity and between-population differentiation is geographic isolation, which enhances genetic drift, thereby lowering within-population genetic diversity and increasing between-population differentiation. This could explain the very low internal diversity of some AA groups (Figure 2A-2C) and high differentiation from other groups, e.g. some groups of Htin (TN1) and Lawa (Figure 4A and 4B; Figure S3) that live in mountainous, isolated parts of northern Thailand. The Lawa furthermore favor intra-marriage (Nahhas 2007), which would also reduce genetic variation in this group.

Moreover, while the expected difference between patrilocal and matrilocal groups holds in some regions (Oota et al. 2001; Besaggio et al. 2007), in other regions patterns of mtDNA and MSY variation do not conform to expectations (Kumar et al. 2006; Arias et al. 2018), which is to be expected if other factors are also influencing patterns of genetic variation (Wilkins and Marlowe 2006).

### Genetic variation and origin of the Mon

The Mon groups showed genetic differences from other AA populations but closer relatedness to the TK populations, especially the central Thai, in both MSY and mtDNA (Figure 2A, 2B, 3A and 3B). Our previous simulation results, based on mtDNA, also supported admixture among the Mon and central Thai groups (Kutanan et al. 2018b). In addition, some Mon groups (MSY: MO3, MO5, MO6 and mtDNA: MO2, MO3 and MO4) exhibit genetic affinities with the Karen (Figure 3B), reflecting genetic heterogeneity and contrasting genetic patterns between MSY and mtDNA. Admixture might be an important factor influencing the genetic structure of the lowland AA-speaking Mon. Archaeological evidence indicates that the Dvaravati civilization of the Mon was centered in present-day central Thailand and southern Myanmar, and had expanded to a large part of mainland Southeast Asia during the 6^th^ to 7^th^ century A.D. (Diffloth 1984; Guillou 1999; Saraya 1999). After the intensification of Thai and Burmese kingdoms, the Mon in Myanmar were conquered by the Burmese during the 18^th^ century A.D.; the ethnic Mon in Myanmar are currently concentrated in the Mon and Karen States (Pon Nya 2001). In Thailand, the present-day Mon are distributed in central Thailand and surrounding areas, with some groups living in the North and the Northeast. However, they are not considered to be the descendants of the ancient Mon Dvaravati civilization in Thailand, but rather political refugees that fled from Myanmar to Thailand during the 16^th^ to 19^th^ centuries A.D. (Ocharoen 1998). However, based on linguistic evidence, the remnants of the Dvaravati Mon population are now considered a distinct ethnic group known as the Nyahkur (BO) whose communities are restrict found in hilly areas along the border between central and northeastern Thailand (Diffloth 1984). In contrary to linguistic evidence, the Nyahkur has no shared haplotype or related to any specific Mon groups, indicating their genetic differences. However, Nyahkur show genetic sharing in both MSY and mtDNA with the Khmer groups (Figure 3A) which reflects their previous connection. In addition, the high frequency of MSY haplogroup O2a* and C* (Figure 1), close genetic relationship to many TK and ST speaking groups (Figure 3B) and highest MPD value for MSY (Figure 2C) indicated later extensive gene flow, promoting the paternal difference of Nyahkur from the Mon and also other AA groups.

Previous genetic studies of G6PD mutations reported a high prevalence of the Mahidol type G6PD deficiency in the Mon, Burmese, and Karen, different from Thai, Laotian and Khmer groups exhibiting the Vientiane-type G6PD mutation (Iwai et al. 2001; Matsuoka et al. 2005; Nuchprayoon et al. 2008). Thus both our results and previous studies indicate a close genetic relationship among Mon, Burmese, and Karen in Myanmar, suggesting a common origin or extensive gene flow. Our previous mtDNA study also revealed genetic relations between some Mon groups (MO1 and MO5) and Burmese, with both of them close to some Indian populations, whereas other Mon groups are closer to the Karen groups (MO2, MO3, MO4) (see details in Kutanan et al. 2018b). In general, genetic mixing among Mon, Karen and Myanmar might have happened before the arrival of the Mon to Thailand, whereas mixing among the Mon and central Thai would have occurred after the arrival of the Mon. However, MSY data for the Burmese are limited, and further MSY studies of populations from Myanmar are needed to confirm this scenario.

A connection between Indian groups and the Mon is suggested by South/Central Asian MSY lineages in the Mon, e.g. R*, H*, J*, L* and Q* (Figure 1), consistent with some mtDNA lineages, e.g. W3a1b, M6a1a, M30, M40a1, M45a, and I1b (Kutanan et al. 2017; Kutanan et al. 2018b). Thus, both mtDNA and the MSY indicated contact between the ancestors of the Mon and Indian. Archaeological evidence also suggests Indian influences, e.g. the symbolism on the Dvaravati coin which indicates the importance of royalty, and includes several motifs associated with Indian precedents of the 1^st^-4^th^ century A.D (Higham and Thosarat 2012).

### Demographic changes

Demographic expansion of Thai/Lao populations are noticeably detected in both paternal and maternal lineages at the beginning of the Holocene, ∼10 kya (Figure 6). In this period, increasing and more stable temperatures might have facilitated population expansion (Wen et al. 2016). The male *N*_*e*_ increase during the Holocene is primarily driven by the O2a2a* and O2a1c* lineages (Figure 8). The Holocene expansion might thus be related to an expansion of HM paternal lineages, as O2* (O-M122*) is thought to have arisen at the beginning of the Holocene near Tibet (van Driem 2017). According to this hypothesis, the bearers of this haplogroup became the progenitors of the “Yangtzean” or Hmong-Mien paternal lineages, and contributed this lineage to the ancient AA who carried O1b1a1a* or O-M95* by sharing of knowledge about rice agriculture. However, further sequencing of MSY lineages belonging to the HM populations are needed to verify this hypothesis.

During the Neolithic period, other significant expansions are observed in almost all ethnicities and many MSY haplogroups, i.e. O1b1a1a1b*, O1b1a1a1a* and R* (Figure 8). Previously, it was suggested that the demographic expansion pattern in the Neolithic in SEA shows strong expansion dynamics, different characteristics than the Paleolithic expansion, and sex-specific expansion patterns, with earlier expansions in female than in male lineages. (Wen et al. 2016). The expansion signals in our results coincide with the beginning of the SEA Neolithic ∼5 to 4.5 kya, during which farming expanded from China to SEA (Bellwood 2018). The farming technology for food production could support a higher population density than hunting-gathering, as agriculture could produce a more steady food supply, and males could avoid hunting dangerous animals; thus, effective population size would increase (Jobling et al., 2004; Yan et al., 2014). The farmer expansion ∼4 kya was probably related to ancestral AA speaking hill tribes with predominantly O-M95* lineages that knew rice agriculture (van Driem 2017; Lipson et al. 2018; McColl et al. 2018). However, the movement of Neolithic groups from southern China to MSEA probably involved not only AA groups but also TK groups (Bellwood 2018). In our study, a Neolithic expansion signal was observed for the MSY in all studied groups, indicating a large demographic expansion and probable admixture among the ancestors of indigenous southern Chinese groups during the Neolithic period. Haplogroup R1a was previously suggested to show a similar expansion, with paternal population growth during ∼6.5 to 4 kya observed globally (Poznik et al. 2016; Wang et al. 2016).

In addition, we found another significant expansion during the Bronze age ∼2 kya that involves TK speaking populations, reflected by some haplogroups prevalent in the TK, e.g. O1a* (Figure 8). This TK related expansion is consistent with the strong expansion detected in the BSP of Xishuangbanna Dai (Figure 7) and corresponds with the results of a recent ancient DNA study (McColl et al., 2018). The southward expansion of the indigenous southern Chinese TK speakers to MSEA was probably driven by the Han Chinese expansion from the Yellow River basin to southern China during the Qin dynasty, starting ∼2.5 kya (Belwood 2018). The migration and expansion of prehistoric TK groups during the Bronze Age has had a profound influence on the modern Thais and Laotians in term of languages and genes. Nowadays TK languages are mostly concentrated in present-day Thailand and Laos, and the relatively high level of TK genetic homogeneity might be also driven by this recent expansion.

Our previous mtDNA modeling to explore the migration and expansion of prehistoric TK groups during the Bronze Age supported the spread of TK languages via demic diffusion and admixture (Kutanan et al. 2017; Kutanan et al. 2018b). Here, a similar modeling approach for the MSY data found weak support for cultural diffusion of TK languages. Although we built the model based on historical sources, the models did not generate the observed variation (Figure S6 and S7), indicating that the analysed models do not correspond to the real paternal population history. A possible reason for this striking difference between maternal and paternal histories might be warfare. Historically, many areas of Thailand saw frequent warfare involving various TK groups ∼200-500 ya (Penth 2000). As a result, forced migrations were imposed upon the losing side and men were taken captive more often than women because men could be used to strengthen the victors’ armies. This could result in a different history for the TK male vs. female population. More complex demographic models could therefore more accurately capture the paternal history of Thai/Lao populations.

It may be that the MSY sequences do not harbor enough information to distinguish among the different demographic scenarios. However, comparison of genetic differences (*Φ*_*st*_ and corrected pairwise differences) among the groups used in the simulations does support a real contrast in the maternal vs. paternal histories for the major TK groups in each region, and also finds genetic heterogeneity among these major groups. The northern Thai people showed closer genetic relationship with the Dai than AA groups in both mtDNA and MSY, supporting the demic diffusion model, whereas the ethnic Lao are closer to Dai for mtDNA but for MSY they are related to both Dai and AA rather equally, suggesting demic diffusion for the maternal history and admixture for the paternal history. The central Thai MSY sequences could be of AA origin because they are genetically more similar to the AA groups than the Dai, supporting cultural diffusion, but for mtDNA they are related to both Dai and AA rather equally, supporting admixture in central Thailand as found previously (Kutanan et al. 2018b). Overall, these results suggest that the demographic history of Khon Mueang, ethnic Lao and central Thais are different, possibly reflecting either different migration routes or different small TK groups that expanded from China (Higham and Thosarat 2012). In addition, different patterns of admixture for males vs. females could have occurred in ethnic Lao and central Thais. Archaeological and historical evidence indicate that prior to the TK migration, there were existing rich civilizations in the area, e.g. the Dvaravati of the Mon and Chenla of the old Khmer. With the arrival of TK groups, the Mon people were incorporated by intermarriage into Tai society and adopted the increasing dominant Thai language as their own (Higham and Thosarat 2012). Our results suggest that there was variation in the pattern of cultural diffusion/admixture involving males vs. females in different groups in the area of northeastern and central Thailand and Laos.

Finally, another more recent expansion signal was detected in the northern Thai AA-speaking Lawa, involving haplogroups O2a2b* and N* (Figure 6 and 8). Historical evidence indicates that after the arrival of the TK groups in northern Thailand, the native Lawa groups were fragmented and moved to the mountains (Penth 2000), resulting in cultural and geographical isolation. In support of this model of isolation and drift, we note that the most negative Tajima’s D value is observed in the LW3 group, which suggests population expansion after a bottleneck (Figure 2D).

## Conclusion

Several factors, e.g. cultural practices, gene flow, genetic drift and geography have influenced the genetic variation and genetic structure of present-day Thai/Lao populations. Here we compared high-resolution mtDNA and MSY sequences and found contrasts in the maternal and paternal genetic history of various Thai/Lao groups, in particular the hill tribes and the AA and ST speaking groups, as well as significant genetic heterogeneity among samples from the same ethnolinguistic group from different locations (Figure 1 and 4). Finally, this new MSY study from Thai/Lao males provides more insight into the past demographic history in the paternal line and, along with our previous mtDNA studies, is generally in agreement with recent ancient DNA studies in SEA that indicate two demographic expansions from southern China to MSEA, with the first involving the ancestors of AA groups and the second involving TK groups (Lipson et al. 2018; McColl et al. 2018). Overall, the contrasting results for the maternal vs. paternal history of some Thai/Lao groups supports the importance of detailed studies of uniparental markers, as such contrasts would not have been revealed by studying autosomal markers in just a few Thai/Lao groups. Additional ancient DNA studies, coupled with more detailed genome-wide data from present-day populations, will provide a complete reconstruction of the genetic history of this region.

## Material and methods

### Studied populations

Genomic DNA was extracted from blood, buccal swab or saliva of 914 males belonging to 57 populations that were classified into 26 ethnolinguistic groups, as described previously (Kutanan et al. 2017; Kutanan et al. 2018a) (Figure 1; Table S5). Ethical approval for this study was provided by Khon Kaen University, Naruesuan University, and the Ethics Commission of the University of Leipzig Medical Faculty.

### MSY sequences

We prepared genomic libraries for each sample using a double index scheme (Kircher et al. 2012) and enriched the libraries for ∼2.34 mB of the MSY via in-solution hybridization-capture using a previously-designed probe set (Kutanan et al. 2018b) and the Agilent Sure Select system (Agilent, CA, USA); further details on the probe design are provided in Table S6. Sequencing was carried out on the Illumina HiSeq 2500 platform with paired-end reads of 125 bp length. Standard Illumina base-calling was performed using Bustard. Illumina adapters were trimmed and completely overlapping paired sequences were merged using leeHOM (Renaud et al. 2014). De-multiplexing of the pooled sequencing data was done by deML (Renaud et al. 2015). The alignment and post-processing pipeline of the sequencing data was described previously (Kutanan et al. 2018b).

### Statistical analysis

#### Genetic diversity and structure

We combined the 914 newly-generated sequences together with 14 published sequences (Kutanan et al. 2018b) belong to two hunter-gatherer populations from Thailand: Mlabri and Maniq (Table S5). This study thus includes 928 MSY sequences from 59 populations and 28 ethnolinguistic groups of Thailand and Laos. To compare with the MSY data, we selected 1,434 mtDNA sequences from the same populations from our previous studies (Kutanan et al. 2017; Kutanan et al. 2018a; Kutanan et al. 2018b) (Table S5). We used Arlequin 3.5.1.3 (Excoffier and Lischer 2010) for the following analyses: summary statistics of genetic diversity within populations, the matrix of genetic distances (*Φ*_*st*_), Analyses of Molecular Variance (AMOVA), and Mantel tests of the correlation between genetic and geographic distances.

#### Genetic relationships

To investigate the paternal relatedness between populations, we performed a discriminant analysis of principal components (DAPC) (Jombart et al. 2010). We grouped our samples based on population sampled, geographic location and ethnicity (Table S5) before running the analysis for 100,000 iterations using *adegenet* 1.3-1 (Jombart et al. 2008).

A correspondence analysis (CA) based on MSY haplogroup counts was performed using STATISTICA 13.0 (StatSoft, Inc., USA). Haplogroup assignment was performed by yHaplo (Poznik 2016). The R package (R Development Core Team) was used to carry out a nonparametric multidimensional scaling (MDS) analysis (based on *Φ*_*st*_ values of MSY and mtDNA), the MDS heat plot with 5 dimensions, showing per-dimension standardized values between 0 and 1, and heat plots of the *Φ*_*st*_ distance matrix and the matrix of shared haplotypes.

To get a broad picture of population relationships in Asia, we included 552 MSY sequences from Asian groups for comparison We downloaded the published Y chromosome sequencing data from the SGDP data set (https://sharehost.hms.harvard.edu/genetics/reich_lab/sgdp/Y-bams/Y.tar) (Mallick et al. 2016), the 1000 Genomes Project (1000 Genomes Project Consortium et al. 2015) and the study of Poznik et al. (2016). We merged and processed all sequencing data through the same pipeline as the samples in our study (Kutanan et al. 2018b). The resulting variant file was merged with data from previous study (Karmin et al. 2015; http://evolbio.ut.ee/chrY/) using Heffalump v0.2 (https://bitbucket.org/ustenzel/heffalump). We subset the variant file to sites that were overlapping the regions present on our capture bait and to samples that had a major haplogroup that was also present in our data set. These samples were combined with our samples; we then removed variant sites for which < 25% of the samples had genotype information, and samples that had > 25% of all sites with missing genotype information. The resulting data set provides 16,684 variable sites, which was imputed using BEAGLE v4.1 (Browning and Browning 2016). Additional details on these populations are provided in Table S7.

#### Bayesian Skyline Plots

Based on Bayesian Markov Chain Monte Carlo (MCMC) analyses, BEAST 1.8.4 was used to construct Bayesian Skyline Plots (BSP) by ethnicity and by haplogroup (Drummond et al. 2012). To avoid a false detection of bottlenecks stemming from the sample collection procedure (Heller et al., 2013), we pooled all populations within the same ethnicity and ran jModel test 2.1.7 (Darriba et al. 2012) to select the most suitable model for each run during the creation of the input file for BEAST via BEAUTi v1.8.2. We used an MSY mutation rate of 8.71 × 10^-10^ substitutions/bp/year (Helgason et al. 2015), and the BEAST input files were modified by an in-house script to add in the invariant sites found in our dataset. Both strict and log normal relaxed clock models were run for each ethnicity and haplogroup, with marginal likelihood estimation (MLE) (Baele et al. 2012; Baele et al. 2013). After each BEAST run, the Bayes factor was computed from the log marginal likelihood of both models to choose the best-fitting BSP plot. Tracer 1.5.0 was used to check the results. We also performed the BSP of compared populations, i.e. Dai, Kinh, Southern Han, Northern Han and Japanese from published MSY sequences (Poznik et al. 2016). The BSPs by ethnicity based on mtDNA genomes were carried out in a previous study (Kutanan et al. 2018a).

#### Approximate Bayesian Computation

In order to investigate the paternal origin of TK groups in Thailand/Laos and their local histories, we employed 5 datasets (encompassing northern Thailand, central Thailand, and northeastern Thailand and Laos) and compared 3 competing scenarios: demic diffusion (i.e., a migration of people from southern China, who are then the ancestors of present-day Thai/Lao TK people); cultural diffusion (i.e., the Thai ancestors were the native AA groups who shifted languages and culture to TK) and continuous migration (i.e., gene flow between a migrant TK and native AA groups) that were developed based on known historical hypotheses (Figure S6). The immigrant and endogenous scenarios postulated an initial split of AA and Dai populations, with a subsequent tree-like split of the target group from Dai (immigrant) or AA (endogenous) populations. The continuous migration model shared the same demographic history as the immigrant model, but also allowed subsequent bidirectional migration between the newly originated population and the AA population. All of the simulations assumed uniform population sizes, fixed separation times based on historical records, a fixed mutation rate of 8.71 × 10^-10^ substitutions/bp/year (Helgason et al. 2015) and a prior distribution for both effective population sizes and migration rates (Table S3). Finally, due to the uneven sample size between the tested groups, we simulated a number of individuals equal to the lowest sample size among the populations in the model.

We simulated the derived site frequency spectrum (unfolded-SFS) for 2,364,048 loci using the fastsimcoal simulator (Excoffier and Foll 2011) with the flag -s, through the software package ABCtoolbox (Wegmann et al. 2011) and running 50,000 simulations for each model. The observed SFS was calculated with the software 4P (Benazzo et al. 2015). To determine the best performing scenario in each set we employed the model selection procedure ABC-RF (Pudlo et al. 2016), which relies on random forest machine learning methodology (Breiman 2001). This classification algorithm is trained on a reference table of simulations and allows the prediction of the most suitable model at each value of a set of covariates (i.e. the summary statistics). Additional details concerning the ABC-RF analyses are described in our previous study (Kutanan et al. 2018b).

## Acknowledgements

We would like to thank all sample donors, village chief and coordinators, i.e. Sukhum Ruangchai, Khamnikone Sipaseuth, Worasitikulya Taratima, Saksuriya Triyarach, Narongdech Mahasirikul, Supada Khonyoung, Dusit Boonmekam, Tharanat Hin-on, Kantaphon Chueahor, Pittayawat Pittayaporn and Waraporn Hongsaphinan for assistance in collecting samples. We thank Murray Cox, Brigitte Pakendorf and Rasmi Shoocondej for valuable discussion. This study was supported by the Max Planck Institute for Evolutionary Anthropology. WK was also funded by the Thailand Research Fund (Grant number RSA6180058), Khon Kaen University (Grant number 6100100) and KKU’s Thai Visiting Scholar 2018. MSr was funded by Naresuan University (Grant number R2561B029).

## Author contributions

W.K. and M.S. conceived and designed the project; W.K., J.K. and M.Sr. collected samples; W.K. and R.S. generated data; W.K., A.B., S.G., L.A., A.H. and E.M. involved data analyses; W.K. and M.S. wrote the paper with input from all co-authors.

## Supplementary Text

### Genetic relatedness among populations

The MA and MN show large differences from the other populations in the heat plots of *Φ*_*st*_ values (Figure S5). However, in general both MSY and mtDNA results show relatively larger genetic heterogeneity of the AA groups vs. genetic homogeneity of the TK and ST groups (Figure S3 and S5). After excluding these MA and MN as outliers, the first dimension of the plot divides the AA populations into two groups: one is diverged from the TK cloud, i.e. KH, KA, SU, TN, BU, BL, LW1 and LW3 and another is interspersed with the TK groups, i.e. SO, MO, BO, and PL (Figure 4A). The MDS heat plot for the MSY supported the divergence of AA populations and also emphasized the similarity between some AA populations and TK populations (Figure 4C). The ST speaking-Karen populations are close to the AA-speaking Mon in the right side of the plot (Figure 4A). Among the TK-speaking populations, BT2 and IS3 are closer to the AA groups on the left side of the plot while the central Thai (CT1-CT7) and one Khon Mueang group (KM7) are closer to the AA groups (MO, PL and LW2) and Karen on the right side (Figure 4A), in agreement with the MDS heat plot for the MSY (Figure 4C). In the second dimension, the Lawa groups are very differentiated (Figure 4A), in accordance with the AMOVA (Table 1) and heat plot results Figure 4C). The heat plot of MSY *Φ*_*st*_ values supports strong genetic homogeneity in the TK and ST groups and also generally shows non-significant differences between the Mon (all groups) and the TK populations, especially with the CT groups, which are different from the other AA speaking populations (Figure 4C). For the MDS of mtDNA (Figure 4B), the Mon generally showed genetic affinity with the TK groups in the center of the plot, with the exception of MO1, MO5 and MO6, which differ from the other Mon groups, as can be also seen in the MDS plot (Figure 4B) and heat plot (Figure 4D). Moreover, contrasting relationships based on the MSY vs. mtDNA was observed for the SK and SO from northeastern Thailand, and BT2 from central Thailand. The mtDNA differentiation from other populations was stronger than that for the MSY for the SK and SO, while opposite was observed for the BT2 (Figure 4A and 4B).

### Thai MSY haplogroup distribution

Among the 928 MSY sequences from Thailand, there are 92 specific haplogroups. Because some of these are subhaplogroups of other haplogroups, we use the following nomenclature: an asterisk denotes a parent haplogroup and all subhaplogroups, while the lack of an asterisk denotes just that specific haplogroup. O1b* is by far the most frequent haplogroup (51.19%) and is present in all populations except the MN, who have only haplogroup K (Figure 1; Table S1). There are two subclades of O1b*: O1b1a1* or O-PK4* (99.37%), and O1b1a2 or O-Page59 (0.63%). Only O1b1a1* was previously reported to be wide spread in northern Thailand (Brunelli et al., 2017), while O1b1a2 is newly reported here, occurring in CT5, YU2 and PU4 (Table S1). There are several subclades of O1b1a1*; the most frequent (50.54%) is O1b1a1a* or O-M95*, which occurs in almost all populations. However, almost half of the AA groups show a very high frequency of O-M95* (>70%), i.e. KH1-KH2, KA, BU, BL, SU, TN1-TN3, MA and LW3 (Figure 1; Table S1) while only two TK populations, i.e. BT2 (94.44%) and IS3 (72.22%) have a high frequency of O-M95*. It appears that the frequency of O-M95* is a major driver of the patterns in the MDS plot in dimension 1 (Figure 4A): we find that O-M95* is at high frequency in the populations on the left of the plot and gradually decreases to very low frequency in the populations on the right side, e.g. MO2, MO4, MO7, BO, CT4 and CT7 (Figure 4A).

O-M95* has also been reported to be frequent in AA populations from Cambodia (Zhang et al., 2015) and Laos (Cai et al., 2011), but infrequent in populations from southern China (Zhang et al., 2015) and rare elsewhere in MSEA (Trejaut et al., 2014). The CA analysis (based on haplogroup frequencies) also supports the divergence of these AA populations, with many O1b* sublineages, e.g. O1b1a1a1b1a (O-B426) and O1b1a1a1a1a* (O-F2758*) (Figure S1). With a total frequency at 11.10%, O1b1a1a1b1a* is prevalent in the LW3 (88.23%), BL (66.67%), LW1 (60.00%), KA (55.56%), and TN3 (47.06%) populations (Table S1).

O2a* or O-M324* is the second most frequent haplogroup with an overall frequency of 25.86%; this haplogroup has relatively high frequency (>50%) in several populations: MO4 (53.85%), PL (66.67%), CT4 (55.55%), CT6 (55.55%), and KM7 (61.53%); and moderate frequencies in some Mon groups (MO5: 35.71%), some TK groups (KM9 (47.05%), LU3 (41.17%), and SH (44.44%), and all ST speaking Karen (KSK1 (41.67%), KSK2 (37.5%), KPA (36.36%)) (Figure 1, Table S1). Interestingly, we also observe a cline in O2a* frequency in the first dimension of the MDS plot that runs opposite to the O-M95* cline: O2a* is at higher frequency in populations located on the right of the plot and decreases in frequency toward the left side (Figure 4A).

Within O2a*, lineage O2a2b1a1a* or O-F8*, which is equivalent to O-M133*, is the most frequent (13.79% total frequency) and occurs in almost all populations, with fairly high frequencies in PL (50%), KM7 (46.15%), CT4 (38.88%), SH (38.88%), KSK2 (37.50%) and KPA (36.36%) (Table S1). O-M133 has been reported in Han Chinese from Taiwan and Thai from Bangkok (12.73-21.59%) (Trejaut et al., 2014), Northern Han (11.36%), Southern Han (9.61%), Kinh (8.70%), Japanese (9.09%) and Dai (24.44%) (Poznik et al., 2016), but is very rare in Malaysia, Indonesia, the Philippines and South Asia (Trejaut et al., 2014; Poznik et al., 2016).

The last subhaplogroup of O observed in our study is O1a* or O-M119*. With a total frequency of 4.53%, O1a* is prevalent in three TK-speaking populations, i.e. LU2 (23.08%), PU4 (22.22%) and PT (22.22%) and occurs at low frequency in several populations, including central Thai (CT2-CT5), Laotian (LA1-LA2) and Lao Isan (IS1-IS3) (Figure 1; Table S1). O-M119* is thus spread across many TK-speaking groups, and also occurs at high frequency in Austronesian populations (Trejaut et al., 2014), indicating shared genetic lineages between TK and AN speaking groups. O-M119* occurs sporadically in just a few AA groups (Mon, SU and TN2) and at low frequency, in agreement with a previous study of Laos (Cai et al., 2011). The observed O-M119* sequences in the AA groups thus might reflect contact with TK groups.

Overall, the SEA-specific O1b* and O2a* haplogroups (with several sublineages) differentiate our studied populations into at least two main paternal sources, and the frequencies of these two haplogroups correspond to the major differentiation in the MDS plot (Figure 4A). However, there are also several minor non-SEA MSY lineages which promote divergence for some populations, e.g. the Lawa groups. Haplogroup N*, a sister clade of O*, is reported to be distributed in north Asian, Tibeto-Burman, and AA groups in southwestern China (Shi et al., 2013). Only one sublineage (N1c2b2 or N-L665) was found in this study (total frequency of 2.80%) and one third of N-L665 was restricted to LW2, enhancing the divergence of this population. It also sporadically occurs in some AA groups (MO2, MO6 and LW1), Karen (KSK1) and TK groups (THK, LU3, CT1, CT2, KM1, KM2 and LA1) (Table S1).

We also observed some minor haplogroups that are abundant in South/Central Asia (Lippold et al., 2014; Karmin et al., 2015; Poznik et al., 2016), e.g. R*, H*, and J*, occurring at total frequencies of 4.18%, 1.40% and 1.62%, respectively (Figure 1). Haplogroup R* is observed in all Mon groups (41.46% of R) except for MO3, and is at highest frequency in MO2 (33.33%) (Table S1). The same proportion of this haplogroup (41.46% of R*) is also detected in all central Thai groups, except for CT2, and is at high frequency in CT3 (26.32%) and CT7 (27.78%), providing more support for genetic connections between Mon and Central Thais. The remaining proportion (17.02%) of R* was found sporadically (only single samples each) in KH1, SU, YU1, IS1, IS4, BT1 and PU1, probably reflecting recent admixture/gene flow. In agreement with these observations, the CA plots show a correspondence between R1a1a1b (R-Z647) and some central Thai and Mon groups (Figure S1). Haplogroup H* shows a similar haplogroup distribution, i.e. occurring in both some Mon (MO2-MO4 and MO7) and some central Thai (CT1 and CT3-CT5) groups. Haplogroup H* also occurs sporadically in the KH1, YU2, SH and KL groups. Elsewhere, haplogroup H* is found in Burmese and Malayan populations (Karmin et al., 2015) and Central Asian populations (Lippold et al., 2014). Haplogroup J2* is distributed in AA speaking Mon (MO2, MO5, MO6 and MO7), BO, BU, TN3 and TK speaking central Thai (CT5 and CT7) and Lao Isan (IS1 and IS2). Haplogroup J2* has been found in many populations from Central Asia (Lippold et al., 2014). Generally, the haplogroup profile indicates genetic affinities between the Mon and South/Central Asian groups, which is consistent with the MDS plots (Figure 4A) and results from mtDNA analyses (Kutanan et al., 2017; Kutanan et al., 2018b).

The other minor haplogroups observed in this study are D1*, G1b and F. Haplogroup D1* was found with the highest frequency in NY (33.33%) followed by KPA (27.27%), and occurs at lower frequency in a few other groups (Table S1). Subclade D1a1a (D-N1) is prevalent in Tibetan groups and ST-speaking Qiang in southwestern China, and less prevalent in MSEA (Shi et al., 2008; Wang et al., 2013). Haplogroup G1b (G-L835/L830) was restricted to the Karen, where it was found in three of the four Karen groups. G1* was previously reported to occur in Southwest and Central Asia (Balanovsky et al., 2015). The CA analysis also supports the divergence of the Karen (KSK1, KSK2 and KPW) based on G1b, the divergence of SK based on F, and the differentiation of NY and KPA based on D1a1a (Figure S1). In general, the occurrence of both SEA specific haplogroups and haplogroups prevalent in North Asia/Tibet and Southwest Asia in the Karen suggest multiple parental sources, in agreement with previous studies based on mtDNA (Kutanan et al., 2014; Kutanan et al., 2018b). Haplogroup F (F-M89) was distributed at low frequency in SO, SK and PU4, who migrated from Vietnam during historical times (Schliesinger, 2000) and was also reported in one Kinh sample from Vietnam (Poznik et al., 2016) and five Lahu samples from southern China (Lippold et al., 2014). The origins of this haplogroup are uncertain but it might have originated in the area of present-day Vietnam and southern China; additional studies of Vietnamese populations would be informative.

## Supplementary Figures

**Figure S1.**
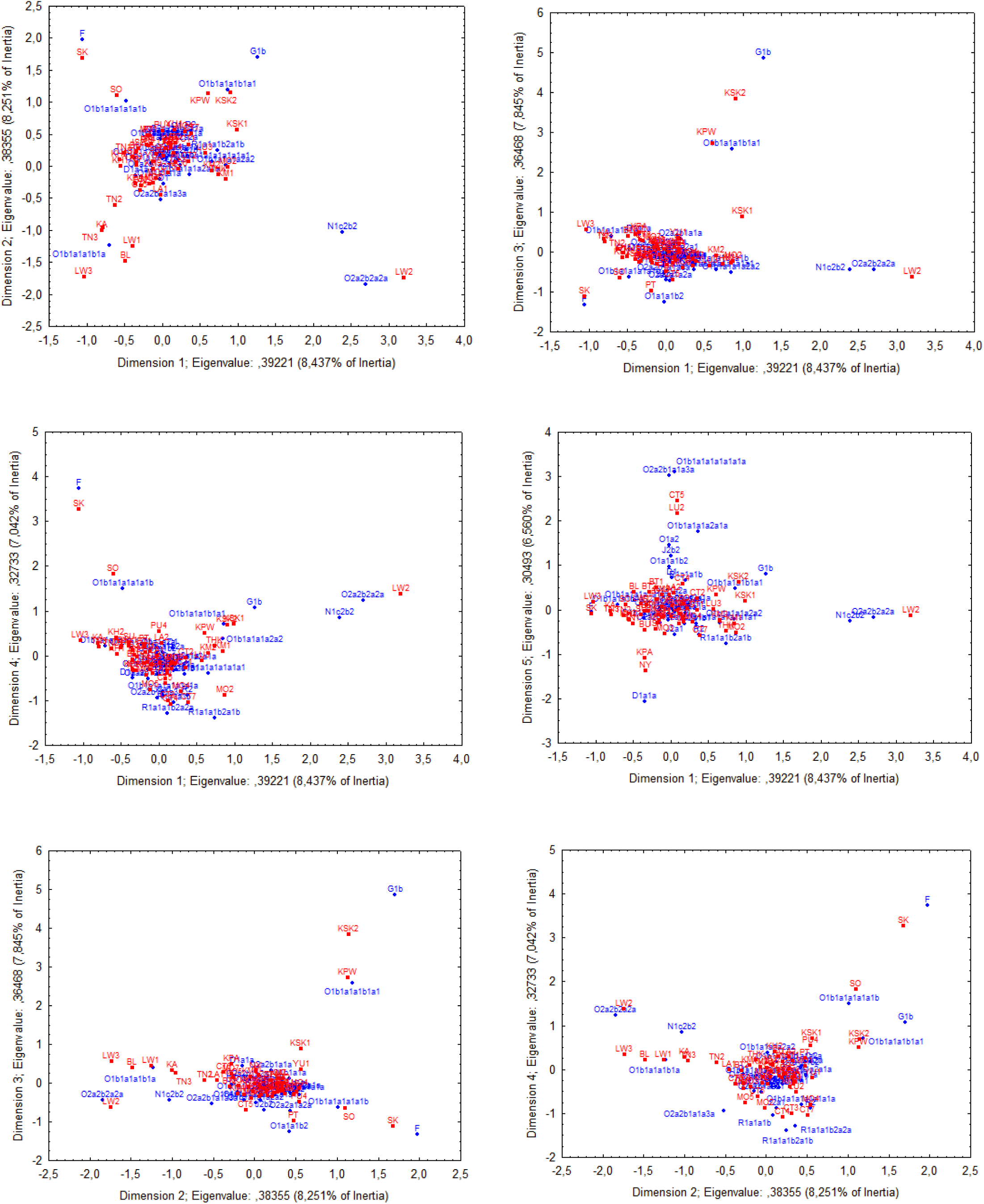
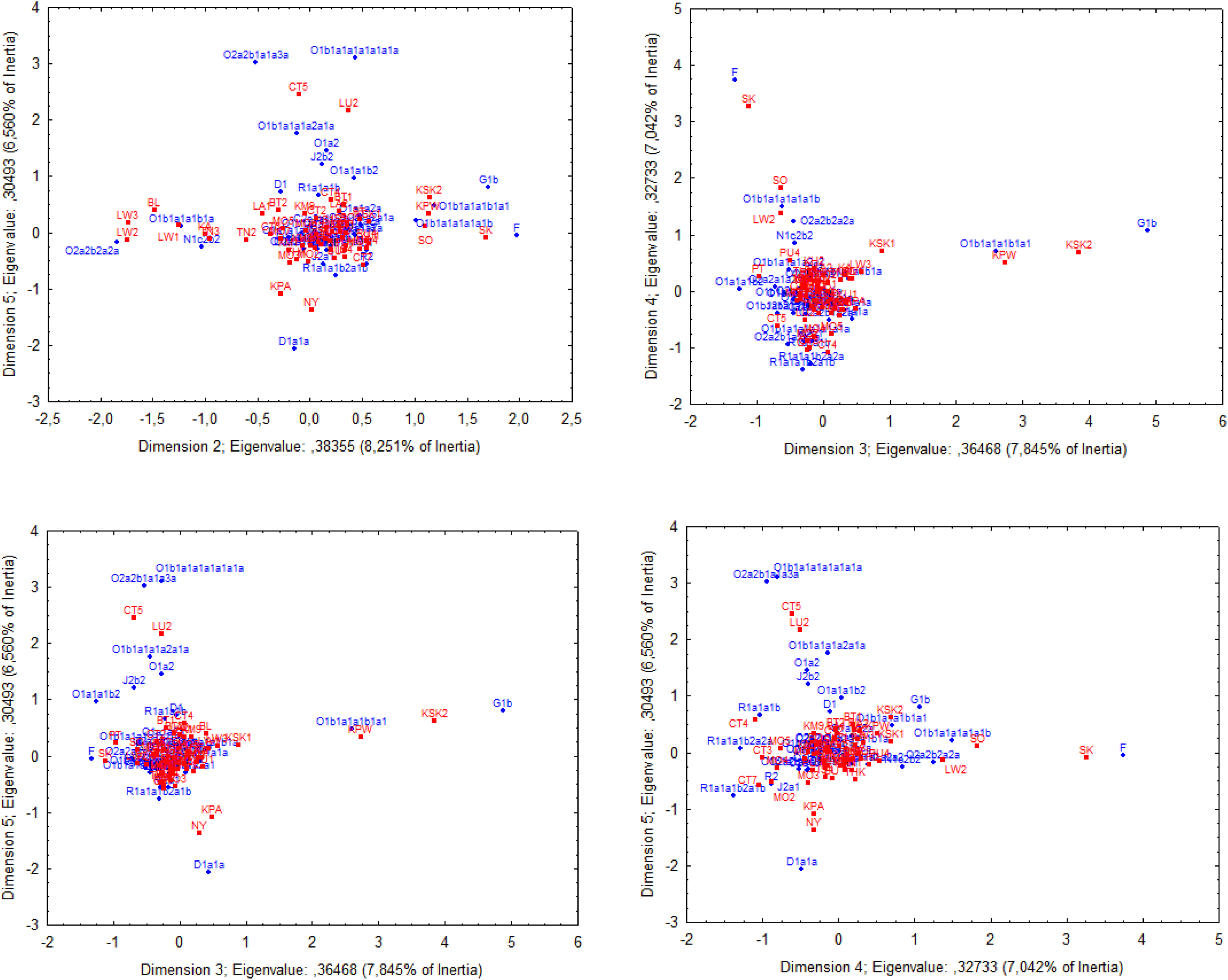
Correspondence Analysis (CA) results based on haplogroup frequency of 58 populations excluding Maniq (MN). Population abbreviations are shown in Figure 1 and Table S5.

**Figure S2.**
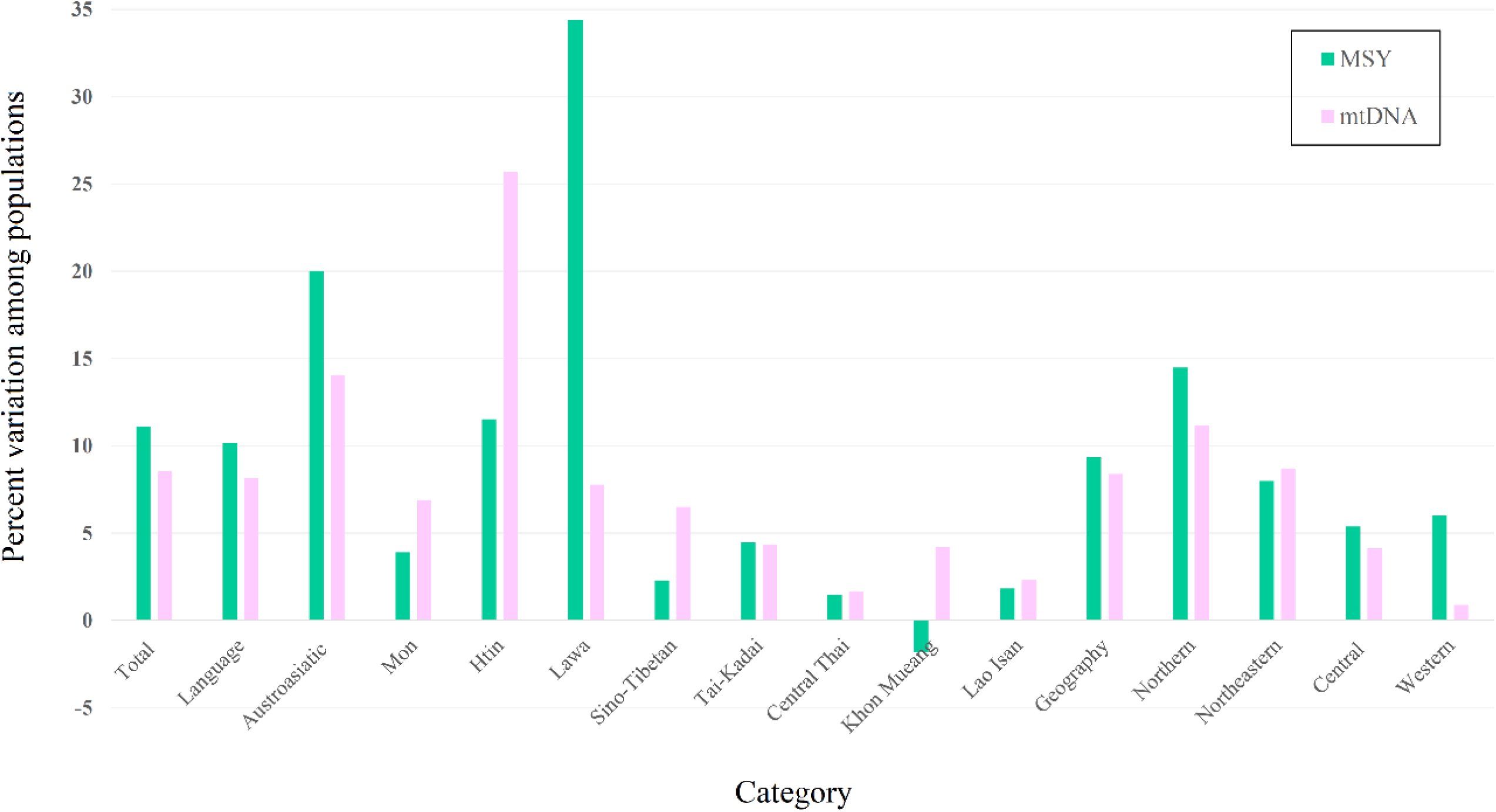
Percent variation among populations in various linguistic or geographic categories, calculated by AMOVA.

**Figure S3.**
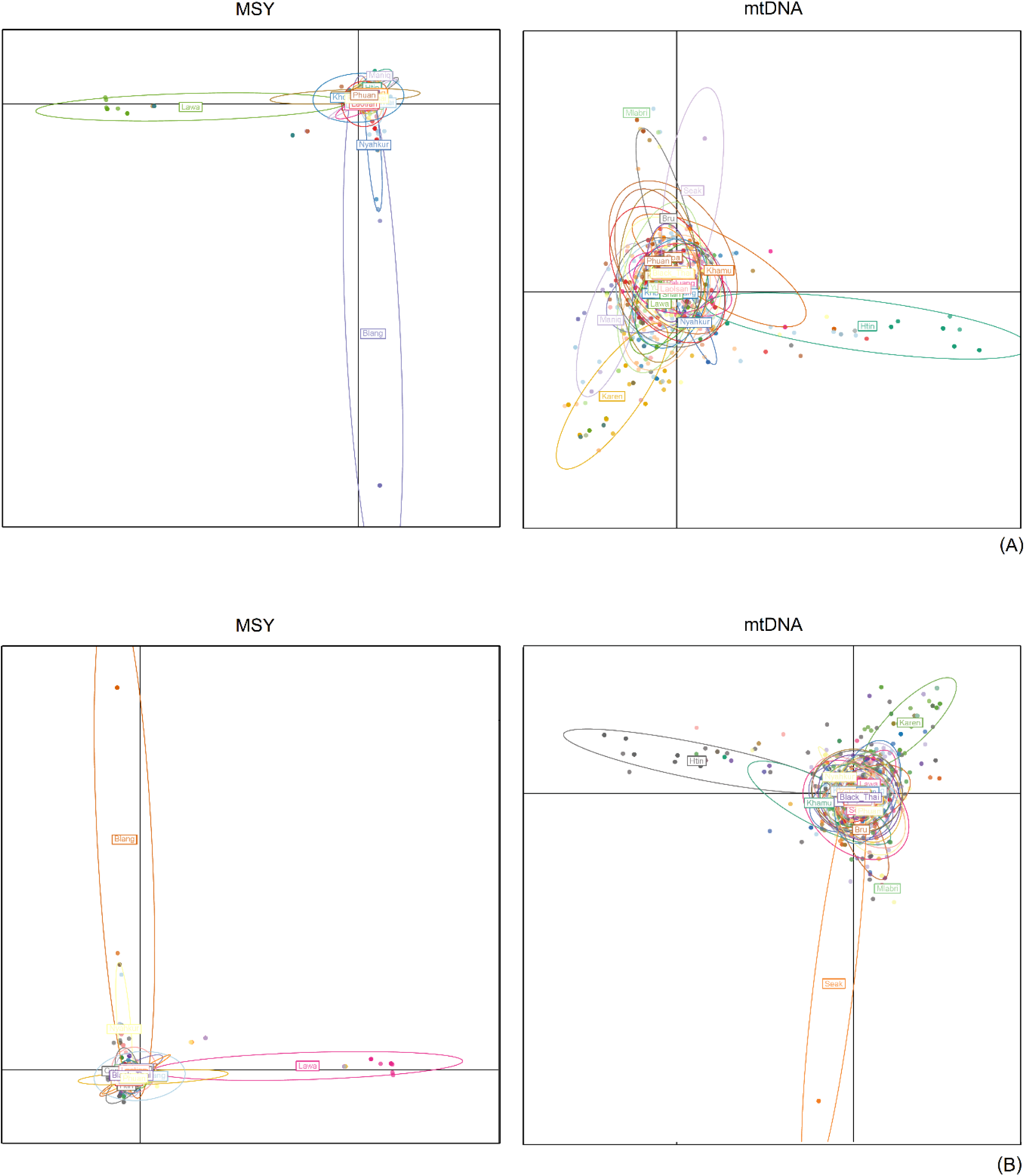
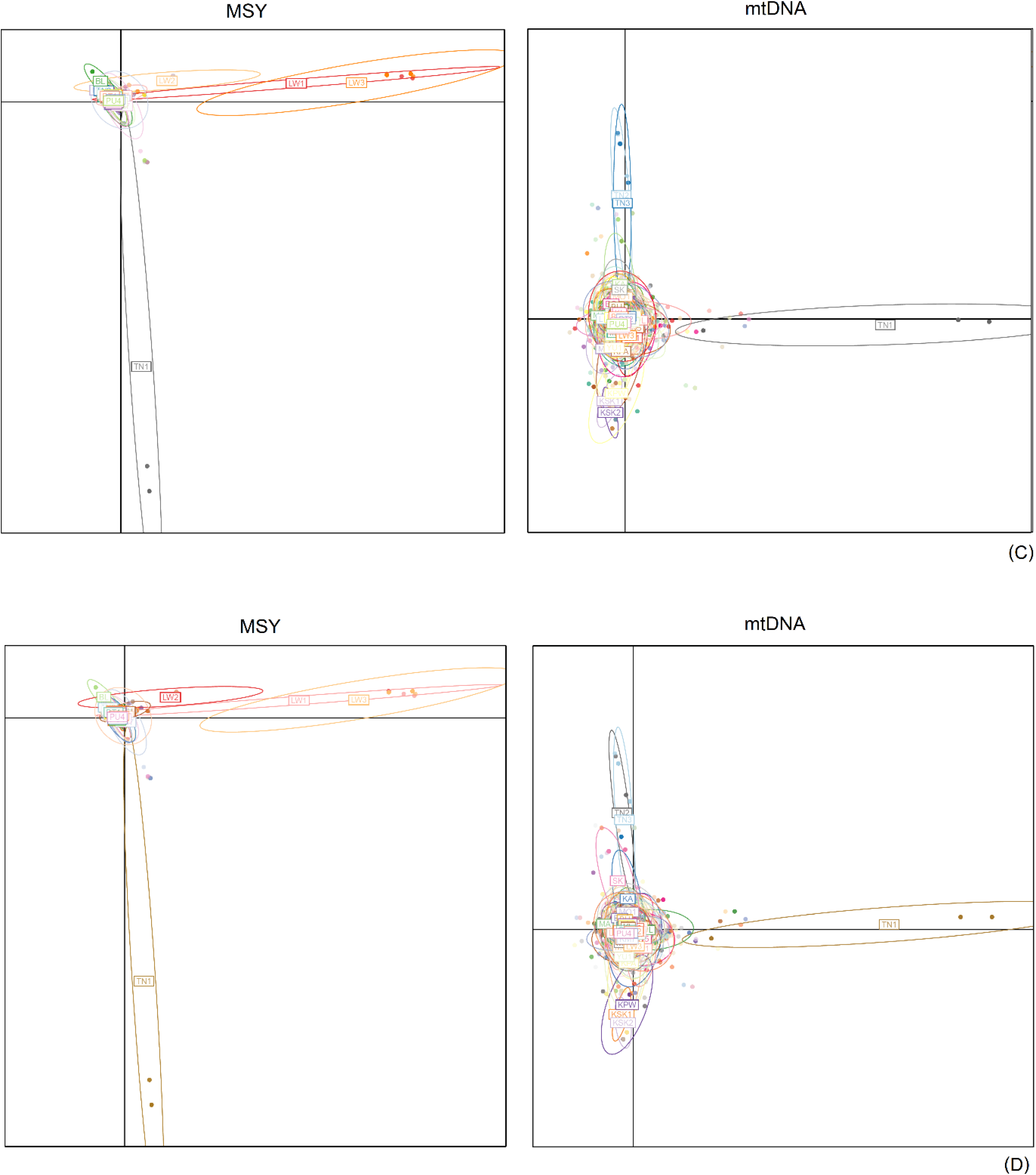
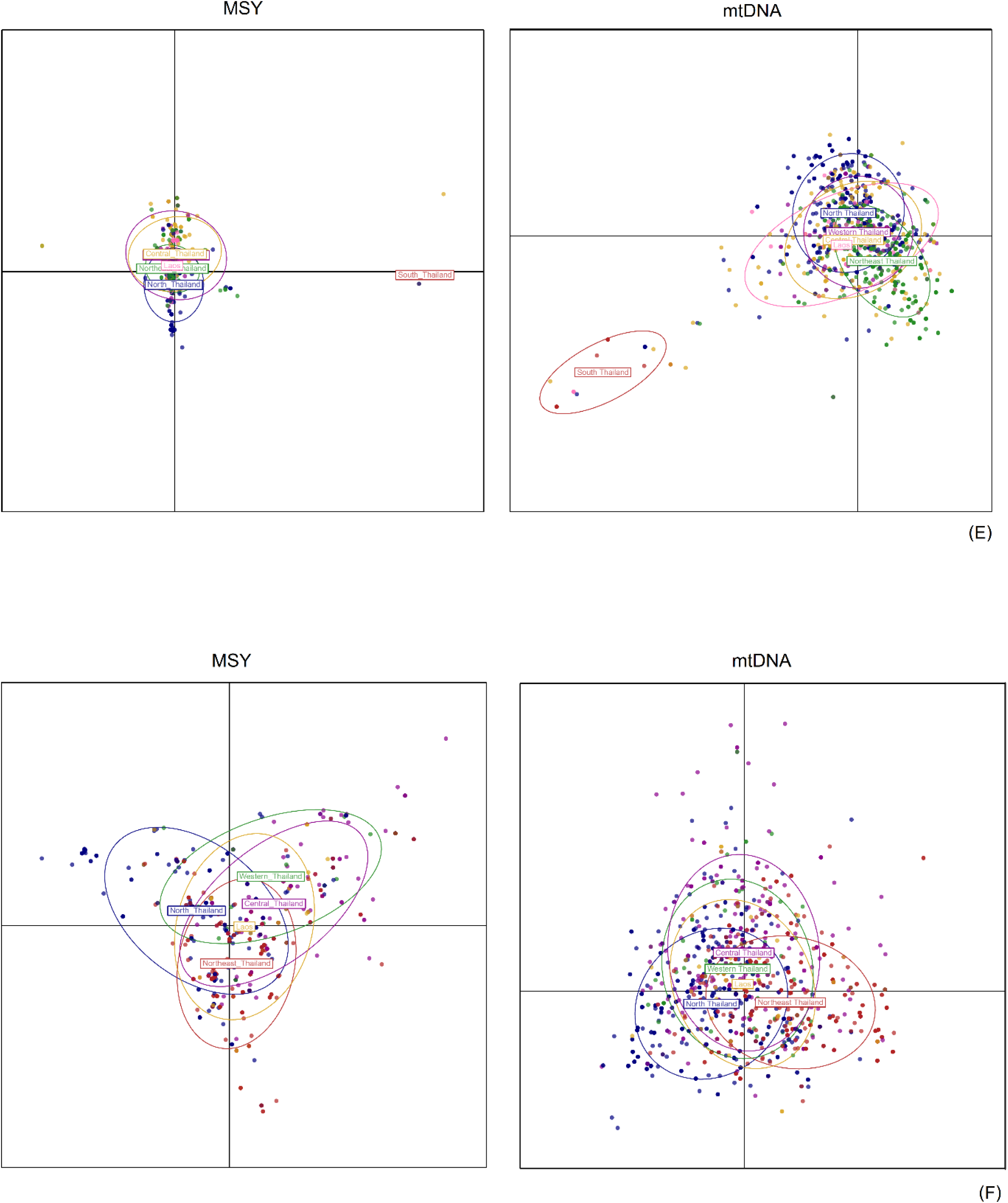
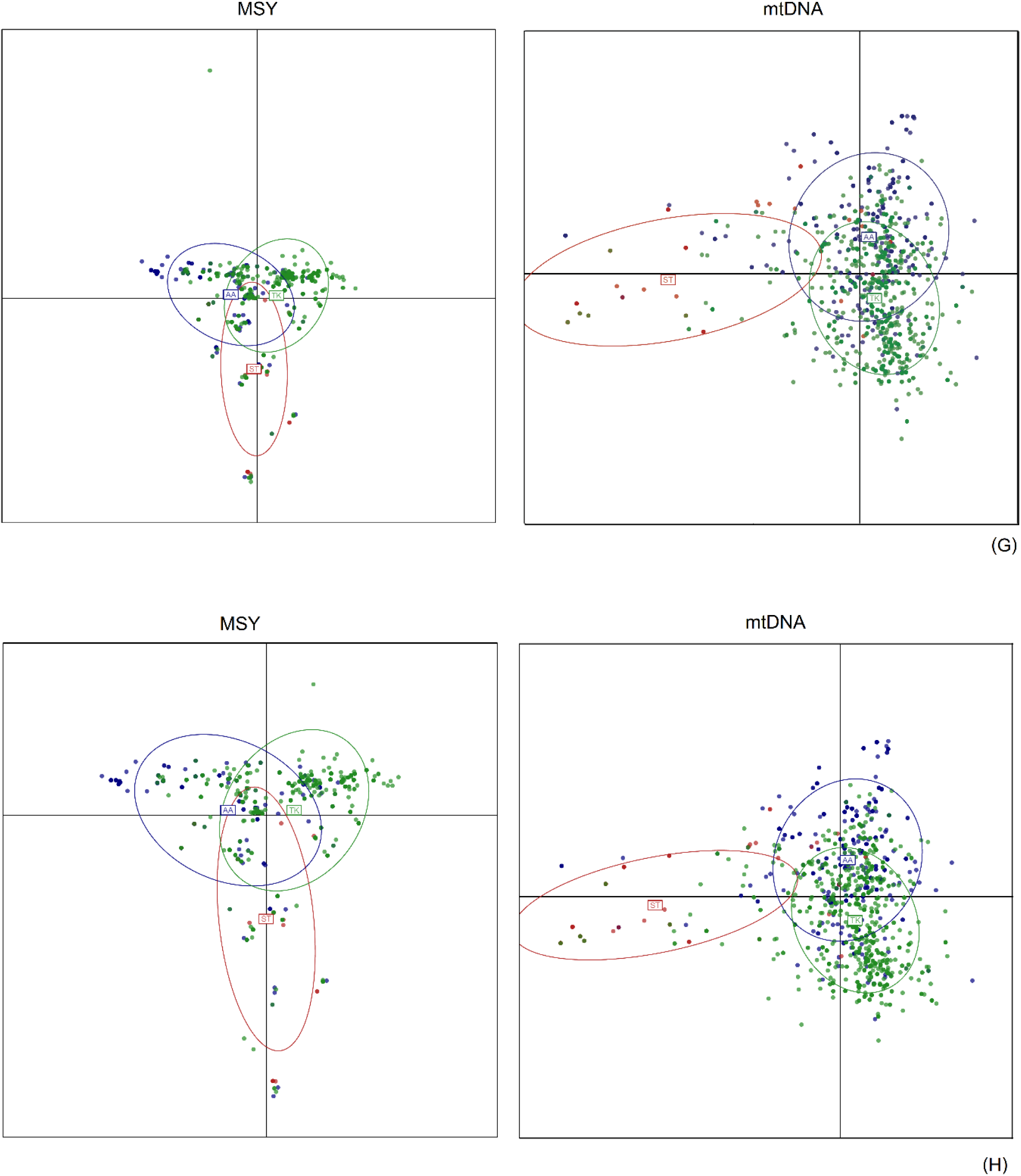
The DAPC results based on ethnicity, population, geography and language (A, C, E and G, respectively). The DAPC results, excluding the Maniq based on ethnicity, population, geography and language (B, D, F and H).

**Figure S4.**
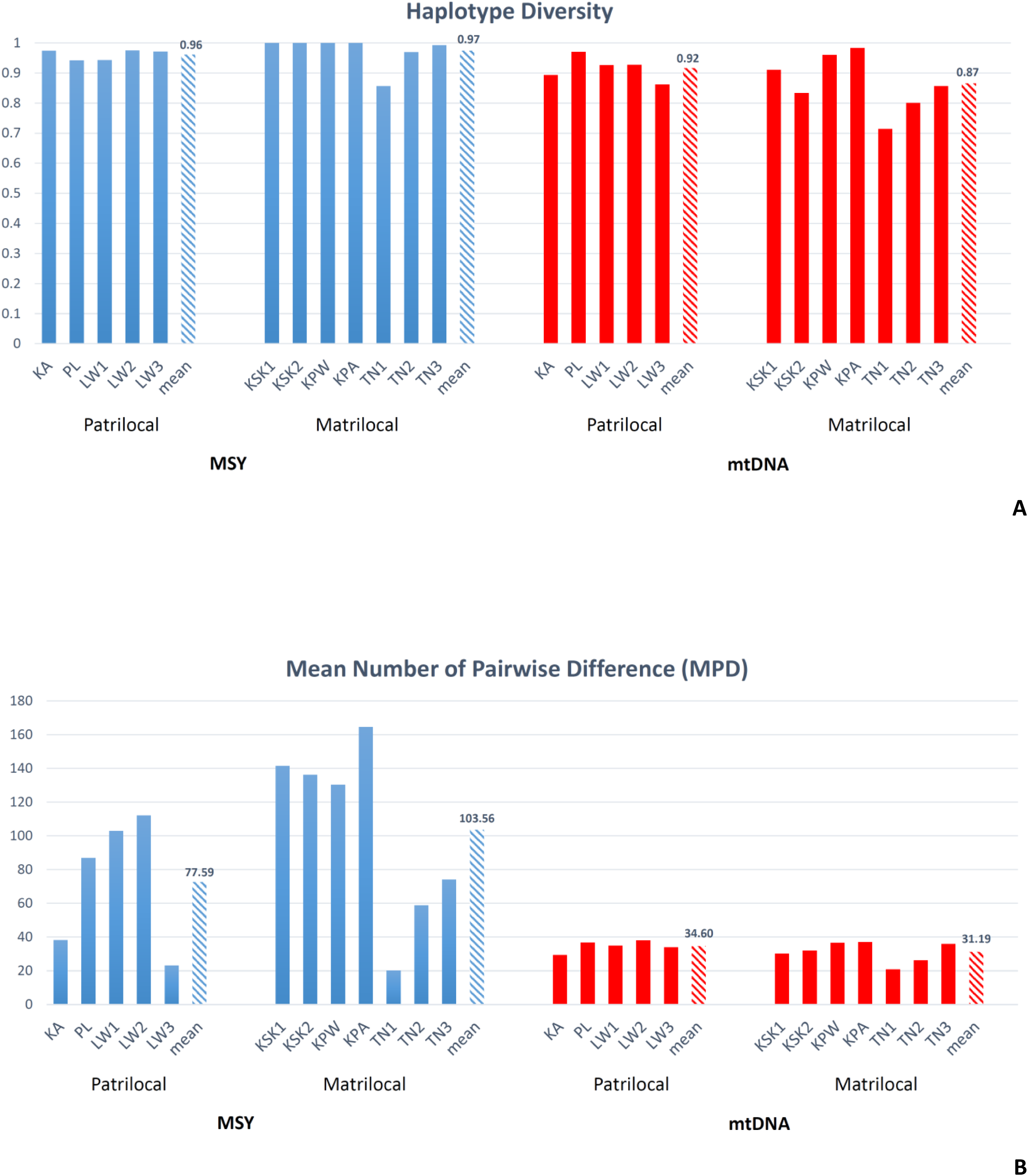
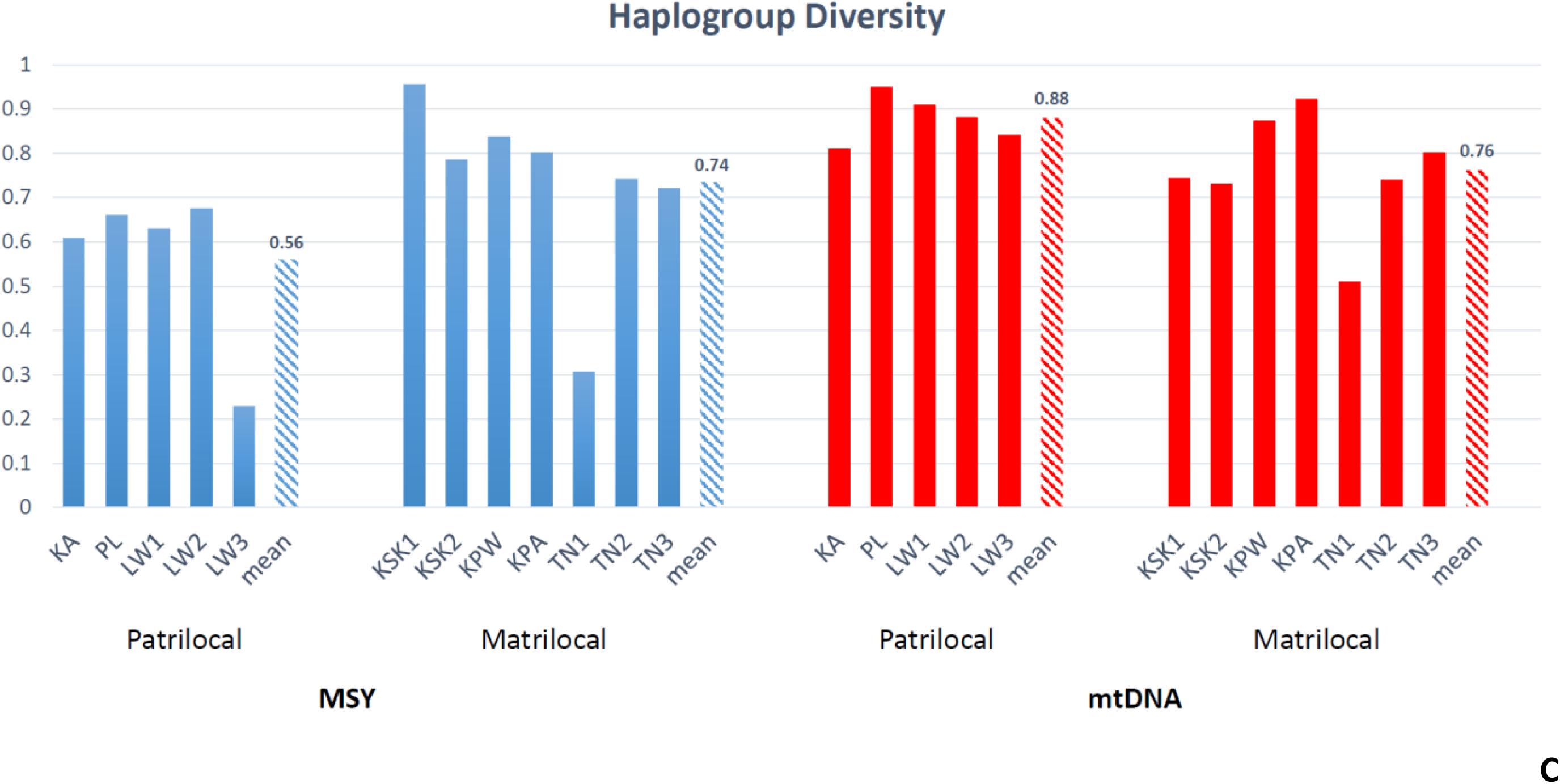
The bar plot graphs of within population genetic variation values, i.e. haplotype diversity (A), MPD (B) and haplogroup diversity (C) in patrilocal and matrilocal groups. The shaded bar in each group indicates the mean diversity in each group.

**Figure S5.**
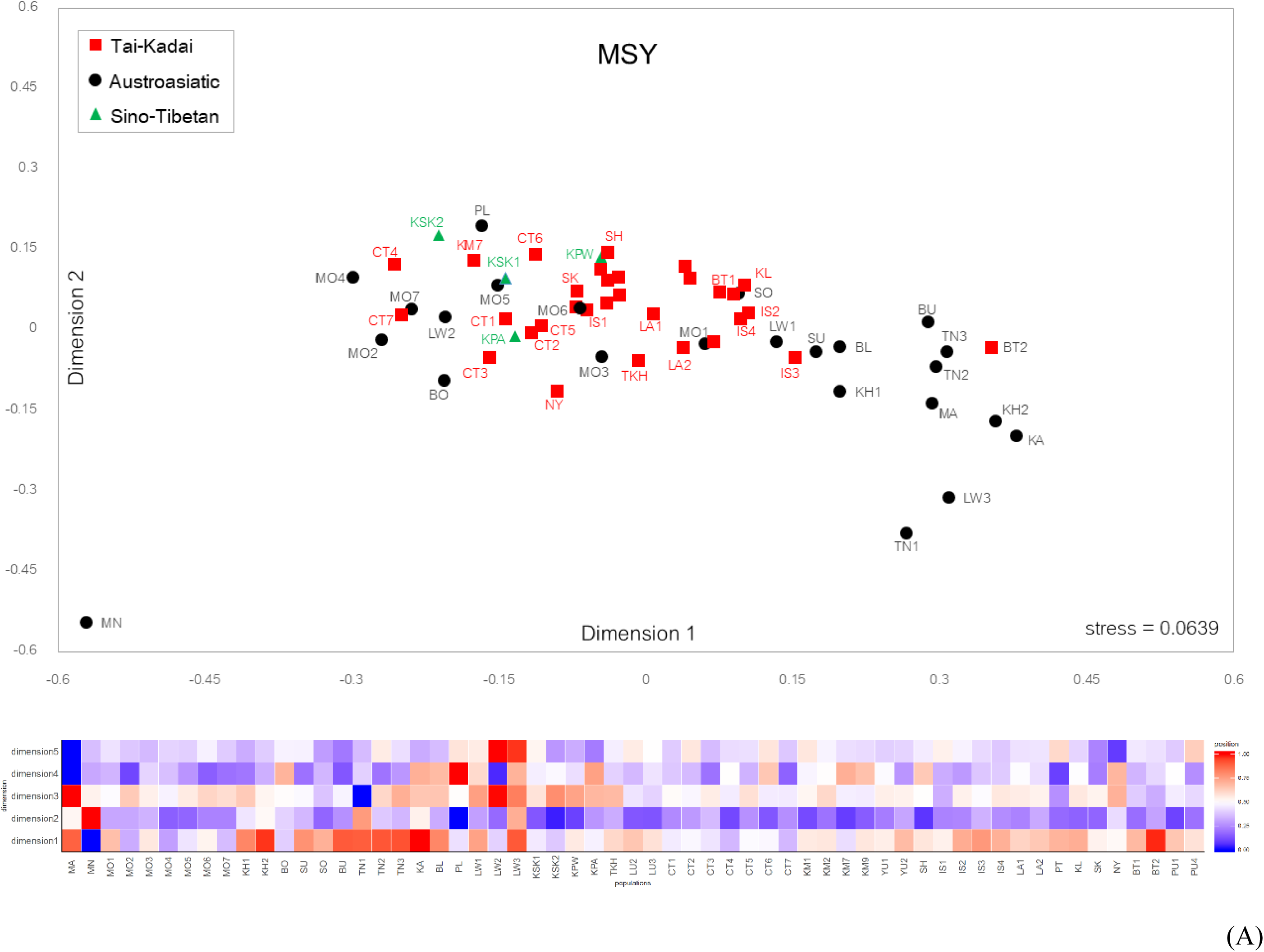
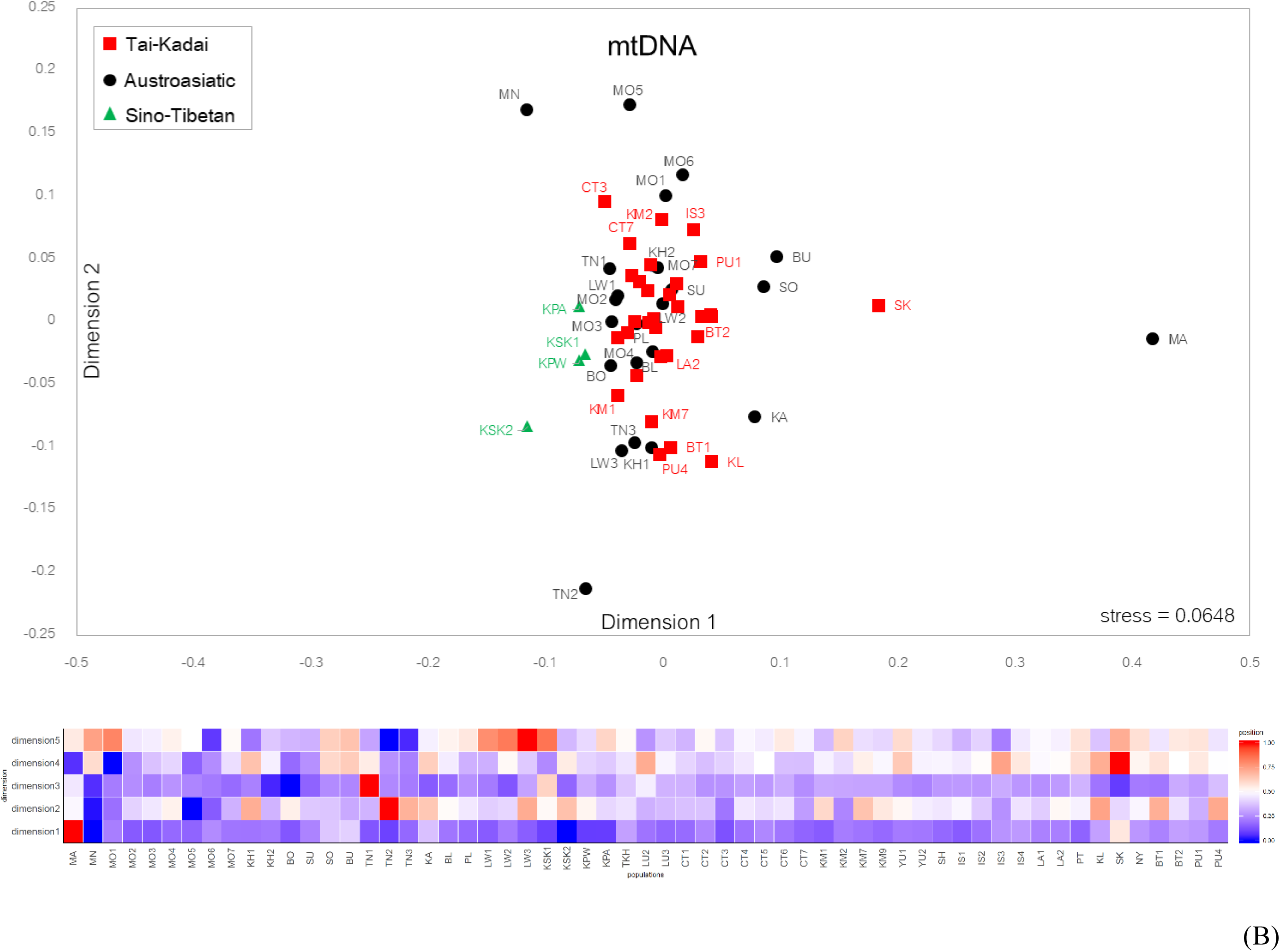
The MDS plot and associated heat plot based on the *Φst* distance matrix calculated from the dataset for 59 populations, for the MSY (A) and mtDNA (B). Population abbreviations are in Figure 1 and Table S5.

**Figure S6.**
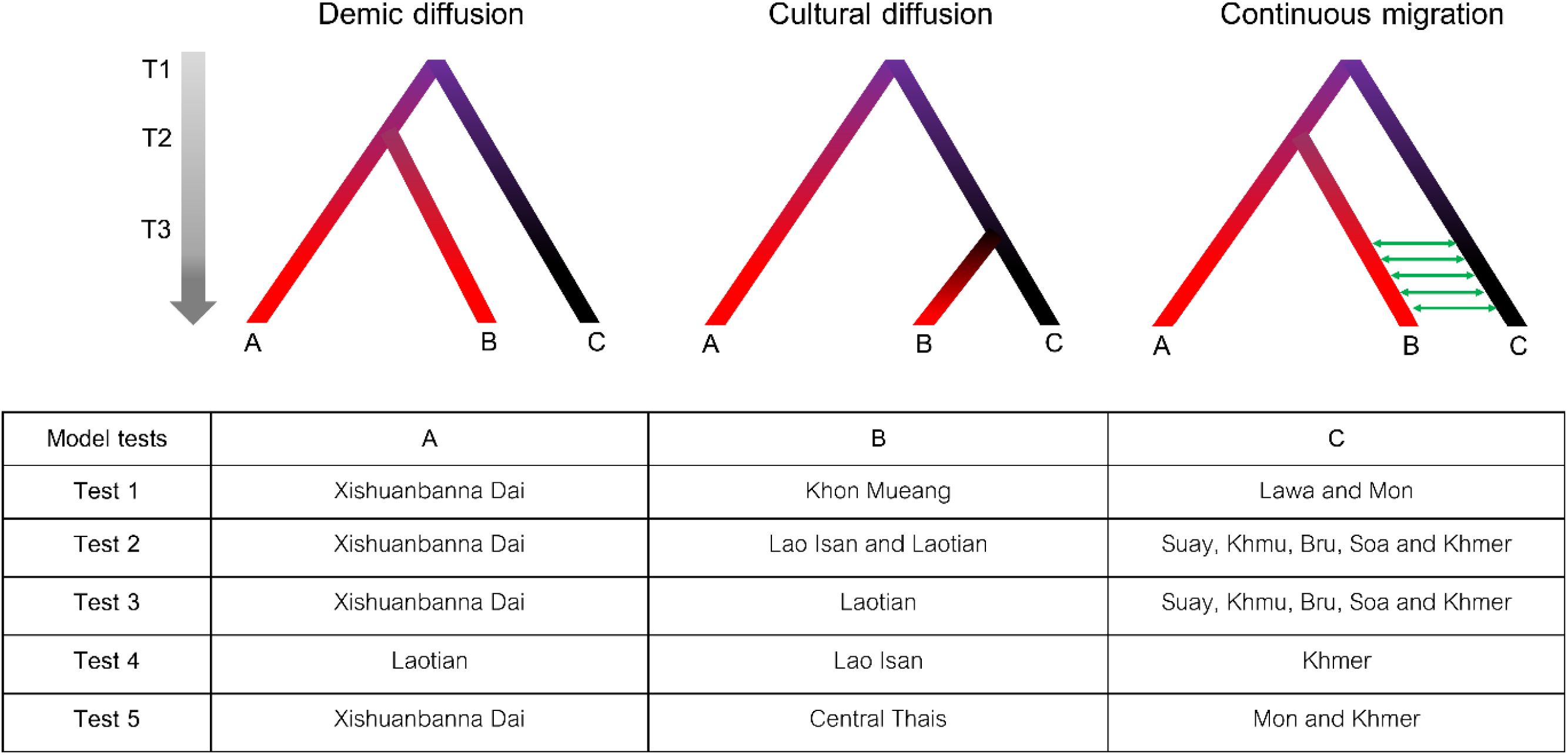
Three demographic models for the ABC analysis (demic diffusion, cultural diffusion and continuous migration). A, B and C represent the different populations and Test 1-5 are the different datasets used in each test. T1, T2 and T3 are either divergence time or time of gene flow.

**Figure S7.**
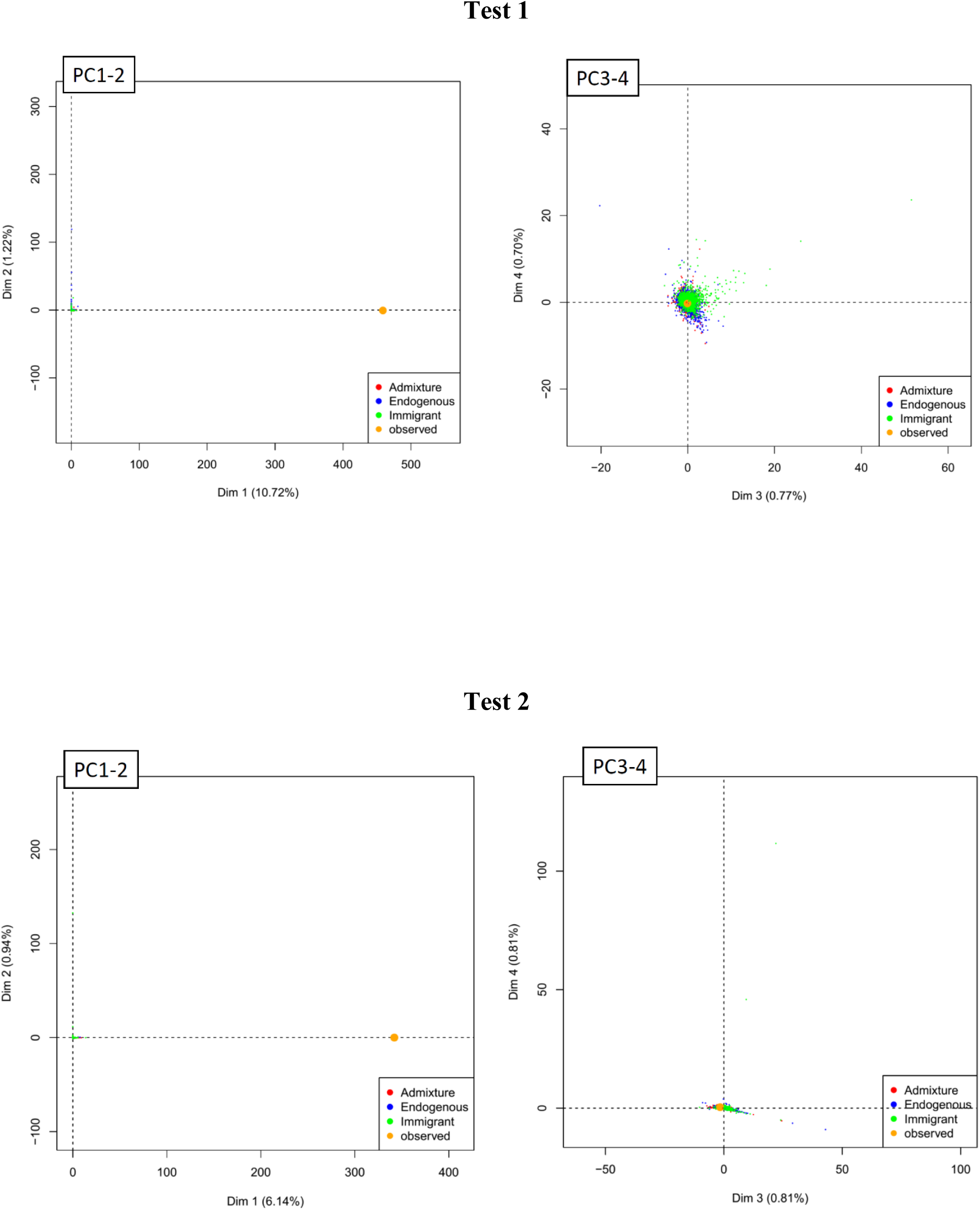
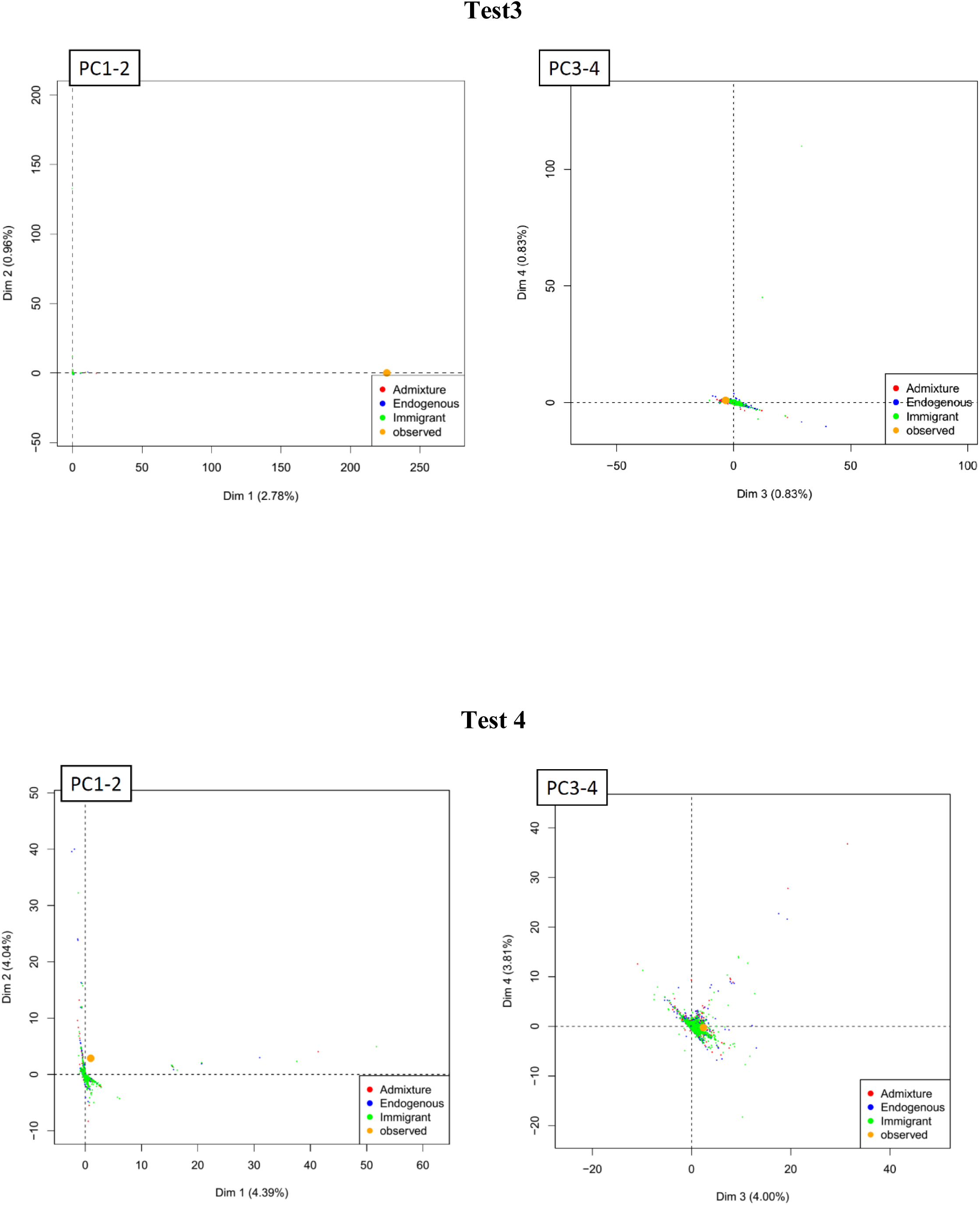
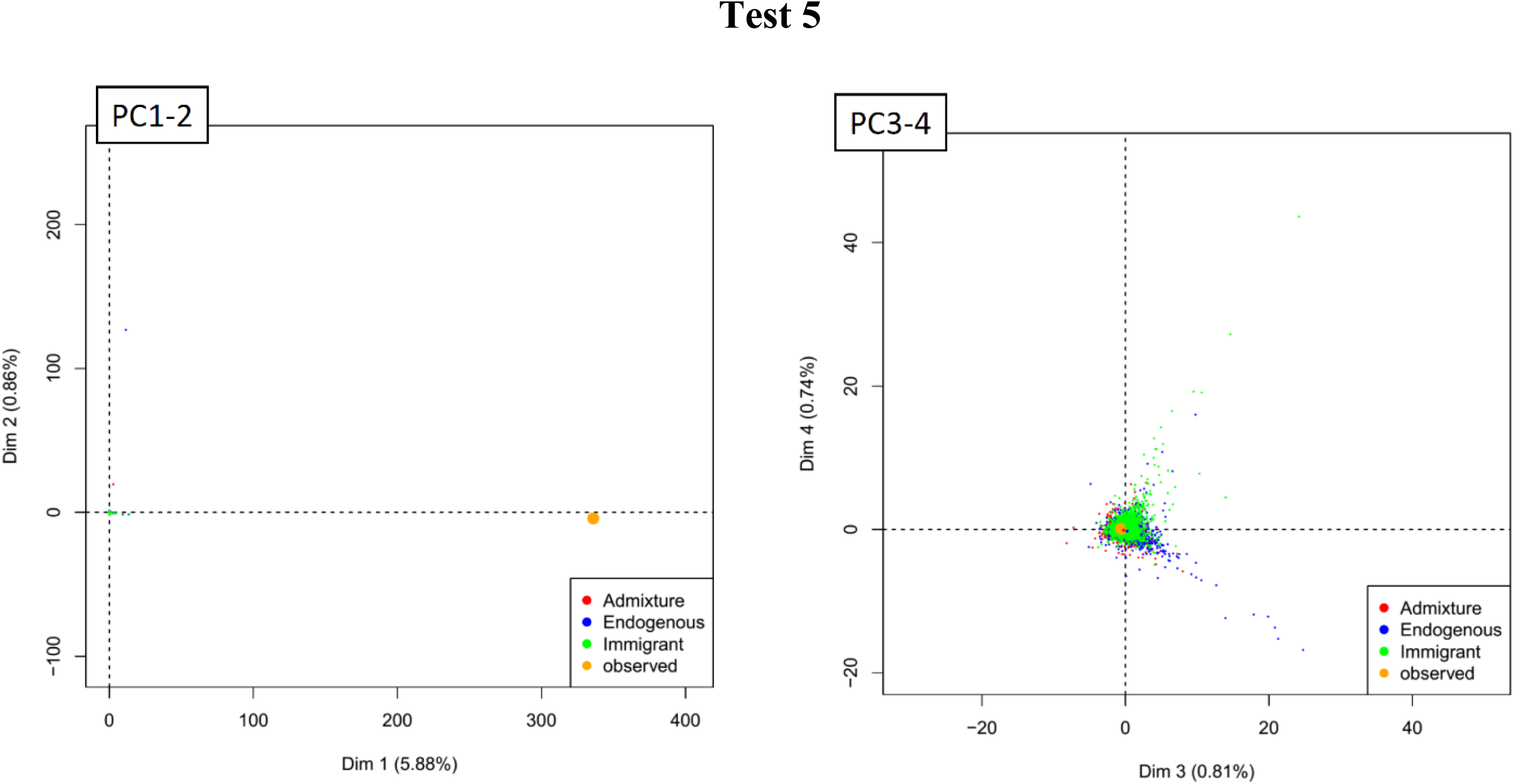
PCA (Principle Component Analysis) analysis based on Dimension 1 and 2 and Dimension 3 and 4 for the fit between the observed data and the simulated data generated by each model for the origin of Northern Thai (Test 1), Laotian and Lao Isan (Test 2), Laotian (Test 3), Lao Isan (Test 4) and Central Thais (Test 5).

**Figure S8.**
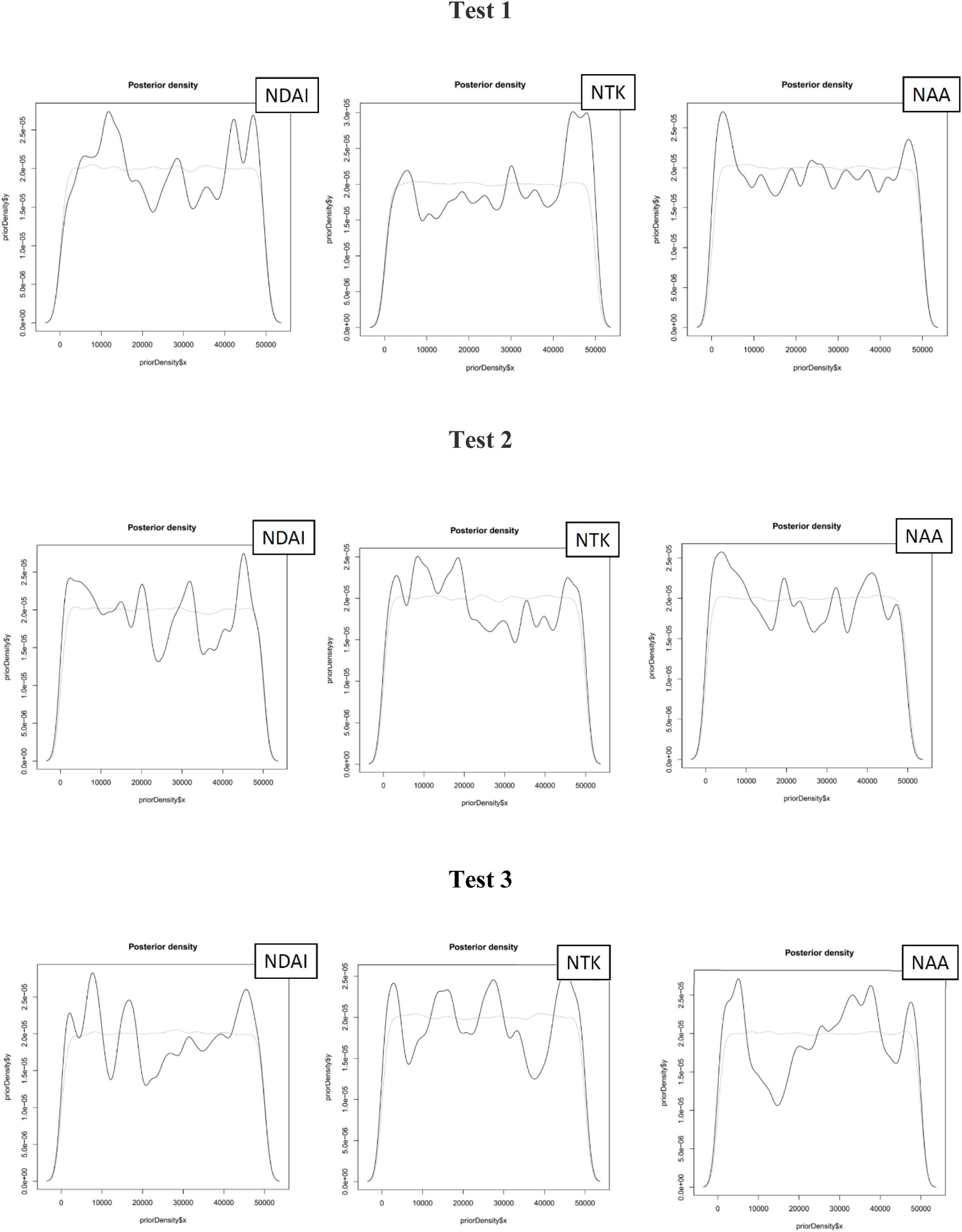
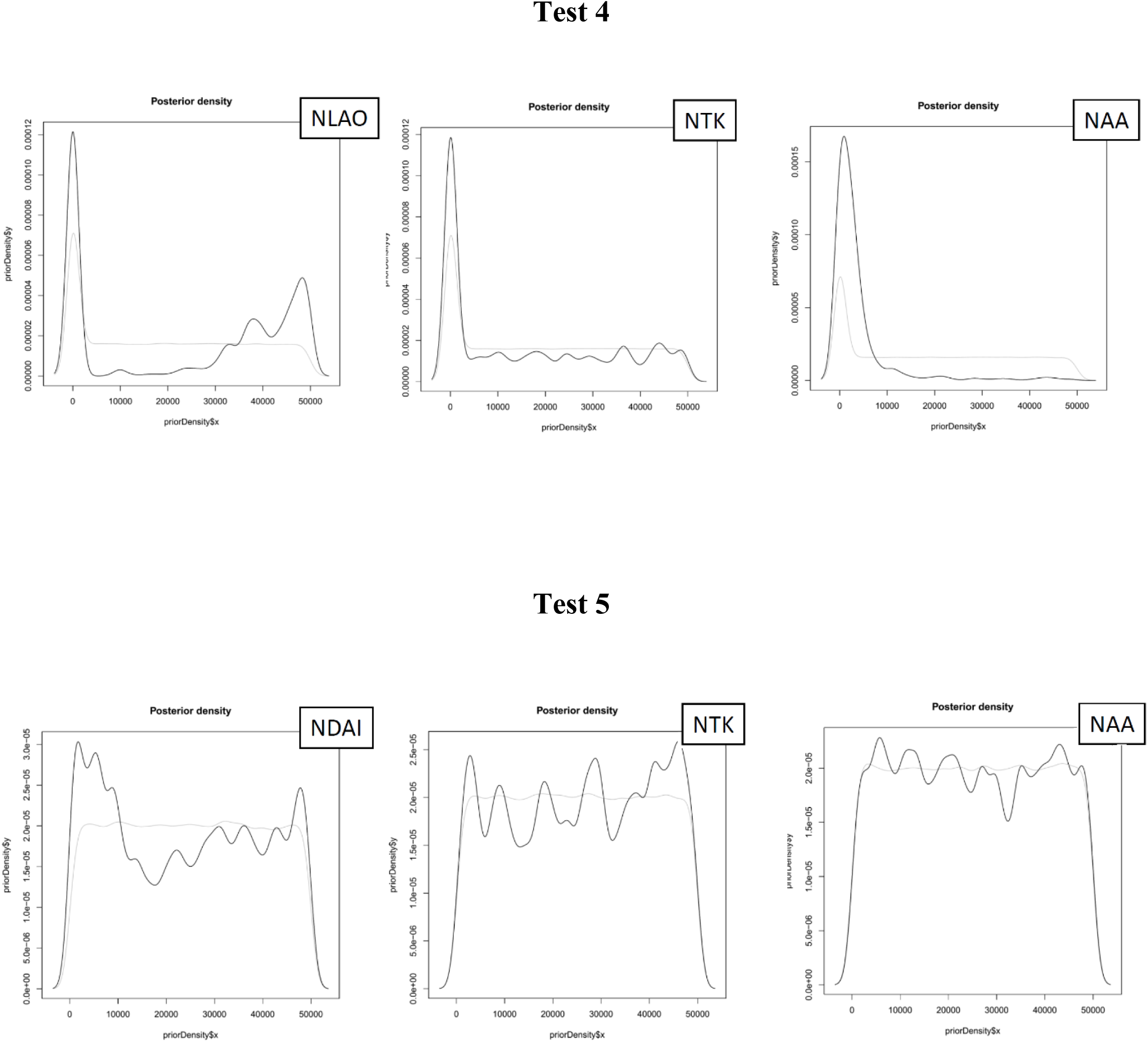
Graphs representing the posterior distribution of each estimated effective population sizes over the extent of the prior range (solid black) and the prior distribution of each parameter (light gray) in each model for the origin of Northern Thai (Test 1), Laotian and Lao Isan (Test 2), Laotian (Test 3), Lao Isan (Test 4) and Central Thais (Test 5).

## Supplementary Tables (excel file)

**Table S1** Haplogroup frequency.

**Table S2** Votes assigned to each model by the Random Forest procedure and posterior probability for the selected model in the ABC analysis.

**Table S3** Parameters estimation for the selected model in each ABC analysis tested.

**Table S4** Genetic differences (Φst and corrected pairwise differences) between each groups of population used in ABC testes. Numbers in parentheses indicate P-values.

**Table S5** General information and genetic diversity values of the studied populations.

**Table S6** MSY probe set details.

**Table S7** Details for the compared populations for MSY data.

